# Design of an *Arabidopsis thaliana* reporter line to detect heat-sensing and signaling mutants

**DOI:** 10.1101/2023.03.26.534276

**Authors:** Anthony Guihur, Baptiste Bourgine, Mathieu E. Rebeaud, Pierre Goloubinoff

**Affiliations:** Department of Plant Molecular Biology, Faculty of Biology and Medicine, University of Lausanne, CH-1015 Lausanne, Switzerland; Institute of Physics, School of Basic Sciences, École Polytechnique Fédérale de Lausanne – EPFL, 1015 Lausanne, Switzerland

**Keywords:** global warming, nanoluciferase, D-amino acid oxidase, heat-shock proteins, heat-stress, HSP20, HSP101, HSP17.3b, chaperones, heat-inducible promoter.

## Abstract

**Background:** Global warming is a major challenge for plant survival and growth. Understanding the molecular mechanisms by which higher plants sense and adapt to upsurges in the ambient temperature, is essential for developing strategies to enhance plant tolerance to heat stress. Here, we designed a special heat-responsive *Arabidopsis thaliana* reporter line that allowed an in-depth investigation of the mechanisms underlying the accumulation of protective heat-shock proteins (HSPs) in response to high temperature.

**Methods:** A transgenic *Arabidopsis thaliana* reporter line named “Heat-Inducible Bioluminescence And Toxicity” (HIBAT) was designed to express from a conditional heat-inducible promoter, a fusion gene encoding for nanoluciferase and D-amino acid oxidase, whose expression was found to be toxic only in the presence of D-valine. HIBAT seedlings were exposed to different heat treatments in presence or absence of D-valine and analyzed for survival rate, bioluminescence and HSP gene expression.

**Results:** Whereas at 22°C, HIBAT seedlings grew unaffected by D-valine, and all survived following iterative heat treatments without D-valine, 98% died following heat treatments on D-valine. The HSP17.3B promoter was highly specific to heat, as it remained unresponsive to various plant hormones, Flagellin, H_2_O_2_, osmotic stress and high salt. Confirming that HIBAT does not significantly differ from its Col-0 parent, RNAseq analysis of heat-treated seedlings showed a strong correlation between the two lines. Using HIBAT, a forward genetic screen revealed candidate loss-of-function mutants defective either at accumulating HSPs at high temperature or at repressing HSP accumulation at low, non-heat-shock temperatures.

**Conclusion:** This study adds insights into the molecular mechanisms by which higher plants sense and adapt to rapid elevations of ambient temperatures. HIBAT was a valuable tool to identify *Arabidopsis* mutants defective in the response to high temperature stress. Our findings open new avenues for future research on the regulation of HSP expression and understanding their role in the onset of plant acquired thermotolerance.

## BACKGROUND

Land plants are increasingly challenged by rising mean seasonal temperatures, which lead to more frequent, lengthy, and extreme heat waves, severe droughts, and devastating fires. Excessive heat at the cellular level can negatively impact the structure and function of macromolecules, such as membranes and proteins, which are particularly thermo-labile (Horvath et al. 2012). Heat can cause transient unfolding of proteins, resulting in the formation of biologically inactive aggregates that tend to expose hydrophobic residues and associate with other proteins and membranes, potentially forming harmful membranal pores (Lansbury and Lashuel 2006). Aggregate interactions with membranes trigger the production of reactive oxygen species (ROS) that in turn elicit apoptosis (Holmes et al. 2014). ROS can also cause direct chemical damage to unsaturated lipids in membranes, oxidize proteins and mutate DNA, ultimately leading to cell death (Mhamdi and Van Breusegem 2018).

Understanding the cellular and molecular mechanisms by which land plants, especially crops, can anticipate and respond to excessive temperatures, is crucial to address the agricultural challenges of global warming. During the morning of a warm summer day, plants must detect an impending noxious heat stress, and accumulate various protective heat-shock proteins (HSPs) and thermo- and ROS-protective metabolites, in order to withstand noxious noon temperatures and survive until late afternoon (Guihur et al. 2022a; Guihur et al. 2022b).

In plants, some heat-induced proteins (HSPs) are enzymes catalyzing the production of heat-protective and ROS-quenching metabolites. Whereas heat-accumulated metabolites, such as glycine betaine and trehalose, can stabilize native proteins and render them less thermolabile (Diamant et al. 2001; Jain and Roy 2009), millimolar concentrations of metabolites can also reduce the heat-induced hyperfluidization and the disruption of biological membranes (Péter et al. 2021). In addition, many HSPs are molecular chaperones. Some heat-accumulating chaperones, such as the HSP20s, do not use ATP to reduce the aggregation of heat-labile proteins (Torok et al. 2001; Reinle et al. 2022; Amen et al. 2021). The HSP20s have also been shown to reduce the heat-induced hyperfluidization and the disruption of membranes (Dutta et al. 2019; Jin et al. 2016). Other conserved families of heat-induced core-chaperones, such as HSP70s, HSP100s, and HSP60s, function as ATP-fueled polypeptide unfolding enzymes (Larkindale and Vierling 2008; Guihur et al. 2022a; Veinger et al. 1998; Finka et al. 2016). These enzymes can unfold heat-misfolded proteins and convert them back into native functional proteins, even under elevated temperatures that are highly unfavorable for the native state (Goloubinoff et al. 2018; De Los Rios and Goloubinoff 2016; Llamas et al. 2021; Fauvet et al. 2021; Tiwari et al. 2023; Avellaneda et al. 2020).

In higher plants, cyclic nucleotide-gated channels CNGC2 and CNGC4, which are members of a family of 20 different stress-responsive membrane-embedded ion channels (Jarratt-Barnham et al. 2021), act as primary heat sensors (Saidi et al. 2009; Finka et al. 2012). Similar to the animal heat sensor TRPV1, that was distinguished in 2022 by a Nobel prize to David Julius (Guihur et al. 2022a; Caterina et al. 1997), CNGC2-CNGC4 hetero-tetrameric channels respond to heat-induced increments in the fluidity of the surrounding plasma membrane, by mediating the transient entry of extracellular Ca^2+^. This initiates a specific signal, involving calmodulin and kinases that hyper-phosphorylates heat-shock transcription factor 1 (HSF1) in the cytosol and induce bound inhibitory HSP70s and HSP90s to dissociate from the inactive HSF1 monomers (Morimoto 1998). Subsequently, HSF1 assembles into active trimers that translocate into the nucleus, where they bind to heat-shock elements in the promoter regions of HSP genes. This instructs chromatin remodeling complexes, such as Arp6, to displace associated inhibitory histones like H2AZ, thus derepressing impending transcription (Kumar and Wigge 2010). This process recruits RNA polymerase to actively transcribe the chromatin-unshackled HSP genes (Zhao et al. 2020; Kumar and Wigge 2010), leading to the massive accumulation of thermo-protective HSPs, predominantly in the cytosol (Guihur et al. 2022a; Guihur et al. 2021).

Here, we designed a stable recombinant heat-sensing reporter line in the model higher plant *Arabidopsis thaliana*, as a tool to carry out genetic screens to identify loss-of-heat-sensing and heat-signaling mutants, expected to be unresponsive to high temperatures, and loss-of-HSP gene repression mutants at low temperature, expected to produce HSPs without heat-shock. A stable reporter *Arabidopsis* line called *“*Heat-inducible bioluminescence and toxicity (HIBAT)” was generated, which expresses a fusion gene encoding nanoluciferase (nLUC) and D-amino acid oxidase (DAO) under the control of a recombinant soybean conditional heat-inducible promoter (HSP17.3B). Heat-induced expression of nLUC-DAO produced a strong bioluminescence signal in heat-treated HIBAT plants grown without D-Valine, whereas it killed most heat-treated plants grown on D-Valine. A forward genetic screen conducted in this study identified several candidate mutants with defective heat shock responses (HSRs) and one mutant exhibiting impaired repression of the HSR at low, non-heat-shock temperatures.

## Results

### Isolation and characterization of HIBAT

A 511-nucleotide fragment, from the soybean heat-inducible conditional promoter HSP17.3B (Supplementary Data S1), has been previously used to design a heat-inducible GUS reporter in *Physcomitrium patens* (Saidi et al. 2005). Here, we aimed to generate a heat-inducible reporter line in the vascular plant *Arabidopsis thaliana* that can conditionally express from the same conditional heat-inducible promoter, a fusion gene encoding for nanoluciferase (nLUC) in N-terminal, fused to DAO (Fig S1). The transformed plants were expected to produce following a heat shock a strong and stable bioluminescence in the presence of furimazine (England et al. 2016), and generate deadly toxic reactive oxygen species in the presence of D-valine, thereby selectively killing plants with an effective heat shock response (Gisby et al. 2012).

*Arabidopsis thaliana* Col-0 plants were transformed using the floral dip method (see Materials and Methods) and homozygous T2 transformants were selected for their ability to produce in the presence of added furimazine, a strong luminescence signal following a 1-hour heat-treatment at 38°C. Since a leaky expression of nLUC-DAO on D-valine at 22°C was expected to be toxic, our selection was further narrowed down to transformants that were totally devoid of nLUC expression at 22°C. Among them, HIBAT was selected as it contains a single copy of the transgene located in an intergenic region of the genome, thereby minimizing potential adverse effects on neighboring essential genes. Whole plant DNA sequencing of HIBAT confirmed a single T-DNA insertion with the transgene at position 26180497 on chromosome 1 (Fig S2). The insertion site was 253 nucleotides downstream of a putative DNA polymerase pseudogene (AT1G69590) and 834 nucleotides upstream of AT1G69600, an expressed gene encoding a zinc finger homeodomain protein.

### HIBAT is very similar to Col-0

At non-heat-shock temperatures (22°C), both HIBAT and Col-0 Arabidopsis thaliana exhibited comparable seed germination percentages, root elongation rates, and fresh weight gains, indicating that the transgene did not negatively impact plant growth and development (Fig S3). However, the cotyledons of HIBAT’s seedlings, but not the true leaves, exhibited a mild decrease of acquired thermotolerance and most turned white, compared to the cotyledons of Col-0 seedlings which stayed green (Fig S4). We nevertheless posited that such a mild heat-sensitive phenotype of HIBAT’s cotyledons, wouldn’t affect our ability to screen for HSR-defective seedlings with 6-8 true leaves. HIBAT was thus retained for its advantageous characteristics of having a single copy insertion of the transgene, in a non-protein-coding region, that while tightly repressing nLUC-DAO expression at 22°C, could drive a massive expression of nLUC-DAO at 36°C.

### Heat-induced expression of nLUC-DAO bioluminescence in *Arabidopsis* seedlings

To assess the heat-inducible expression profile of the nLUC-DAO transgene, HIBAT and Col-0 seedlings underwent two-hour exposure to a range of temperatures between 22°C and 38°C. This was followed by a two-hour period at 22°C to facilitate the completion of HSP synthesis. Seedlings were subsequently treated with a 1/100 furimazine solution. In the absence of prior heat shock, neither HIBAT nor Col-0 seedlings emitted any light. In contrast, following HS, only HIBAT seedlings displayed strong bioluminescence (Fig 1A).

**Figure 1:**
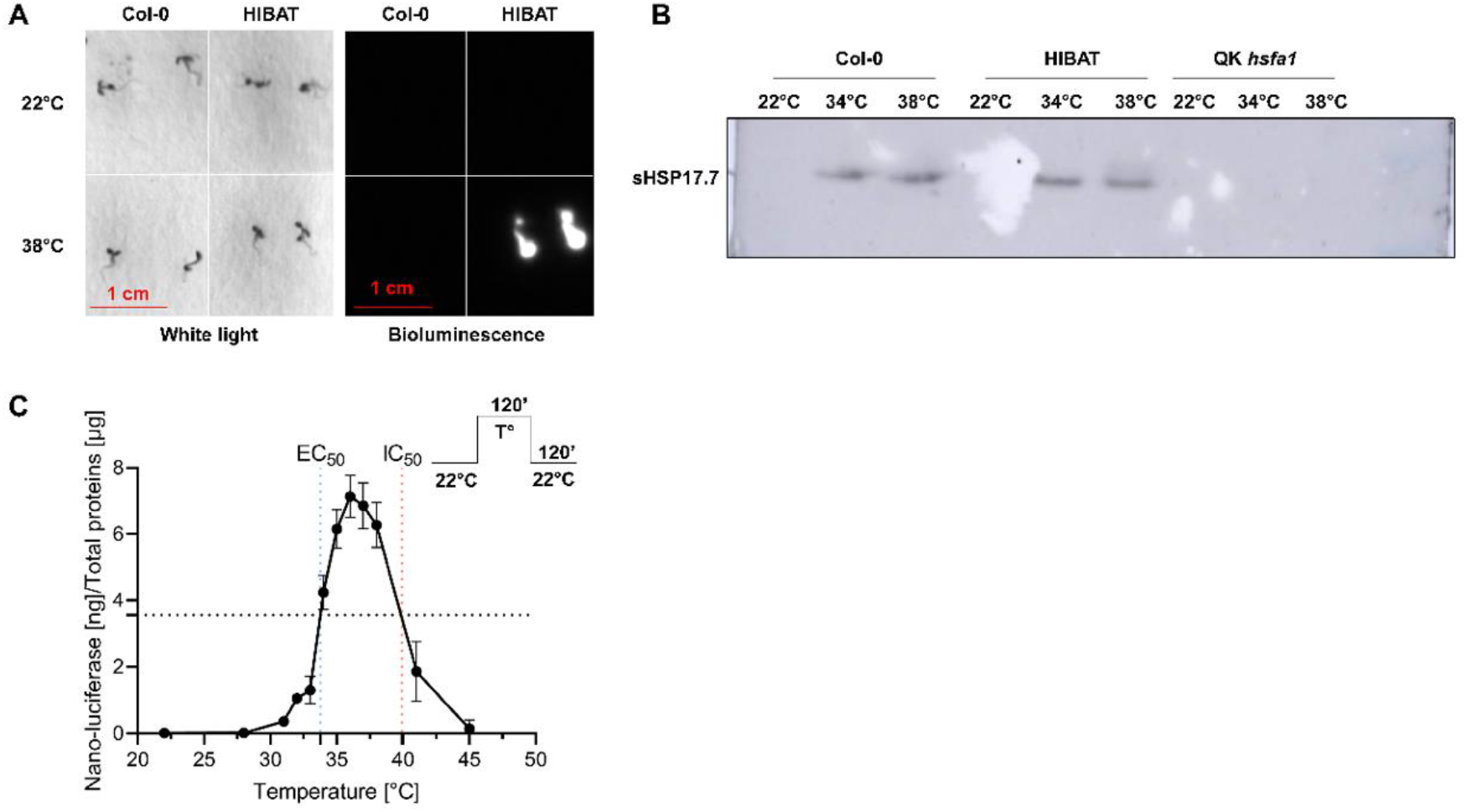
Heat-induced expression of nLUC-DAO bioluminescence. **A**) 2-week-old HIBAT seedlings grown at 22°C were exposed for 2 hours at 36°C and sprayed with furimazine diluted 1/100e. Pictures were taken and following 2 hours recovery at 22°C. **B**) Immunoblot detection of sHSP17.7 in Col-0, HIBAT, and quadruple hsfa1 KO mutant. Leaves were exposed at 22°C, 34°C, or 38°C for 2 hours, then for 2 hours at 22°C. **C**) Expression of the heat-inducible nLUC-DAO. HIBAT seedlings were exposed at indicated temperatures for 2 hours and following 1 hour of recovery at 22°C. The concentration of expressed nLUC-DAO proteins was measured and compared to a calibration curve with purified nLUC (means ± S.D, n = 8)

The light signal’s reliance on HSP17.3B was verified through immunoblots and extended to other endogenous *Arabidopsis* HSPs. At 22°C, endogenous *Arabidopsis* sHSP17.7 and HSP101 were virtually undetectable in both HIBAT and Col-0 seedlings. Following heat shock at 34 and 38°C, both sHSP17.7 and HSP101 strongly accumulated in Col-0 and HIBAT, but not in the quadruple *hsfa1* knockout *Arabidopsis* mutant used here as a negative control (Fig 1B)(Liu et al. 2011). This observation coincided with a substantial accumulation of nLUC-DAO exclusively in HIBAT, as determined by nanoluciferase activity in total protein extracts (Fig 5S). These results demonstrate that the transgene-encoded nanoluciferase activity in HIBAT serves as a faithful reporter for heat-induced expression levels of endogenous HSPs, which are distally encoded from the transgene in the genome.

To determine the temperature-dependent activation profile of the recombinant HSP17.3B promoter, nanoluciferase activity was measured in soluble extracts from HIBAT seedlings following various heat treatments between 22°C and 45°C. Below 28°C, HSP17.3B-mediated expression of nLUC-DAO was tightly repressed. Above 28°C, expression levels strongly increased, with half-maximal expression observed at 33.8°C and a maximal expression at 36°C. Nanoluciferase overexpression declined above 36°C, with an IC_50_ at 40°C (Fig 1C). Since *in vitro*, nanoluciferase is highly thermostable (Tm = 60°C, Promega), this decline is likely due to the heat-impairment of the protein synthesis machinery, rather than to the heat-denaturation of nLUC *per se*.

### Impact of D-Valine on HIBAT survival at 22°C and following heat-treatments

At 22°C, both Col-0 and HIBAT seedlings grew undisturbed by the presence in the growth medium of up to 30 mM D-valine (Fig 2). After two heat treatments, each consisting of 2 hours at 38°C, separated by a 2-hour period at 22°C, the majority of HIBAT seedlings grown on 25 mM D-valine perished within several days, while all wild-type Col-0 seedlings continued to grow normally and thrive. This demonstrates that the expression of nLUC-DAO, which is toxic on D-valine, is tightly repressed in HIBAT at 22°C. Moreover, following heat-treatments without D-valine, which are not lethal *per se* for both Col-0 and HIBAT, the amount of heat-accumulated nLUC-DAO in HIBAT was sufficient to generate on D-valine, enough toxic chemicals (keto-acid-3-methyl-2 oxobutanoate, NH3, and H_2_O_2_) to kill 98% of the seedlings (Fig 2B, Fig S5). Consequently, HIBAT gathered the necessary characteristics for an *Arabidopsis* reporter line to specifically conduct genetic screens aiming at identifying loss-of-function heat-sensing and heat-signaling mutants. These mutants were anticipated to withstand heat treatments in the presence of D-valine, and loss-of-HSP repression mutants were expected to emit low levels of luminescence and HSPs at low temperatures (in the absence of D-valine).

**Figure 2:**
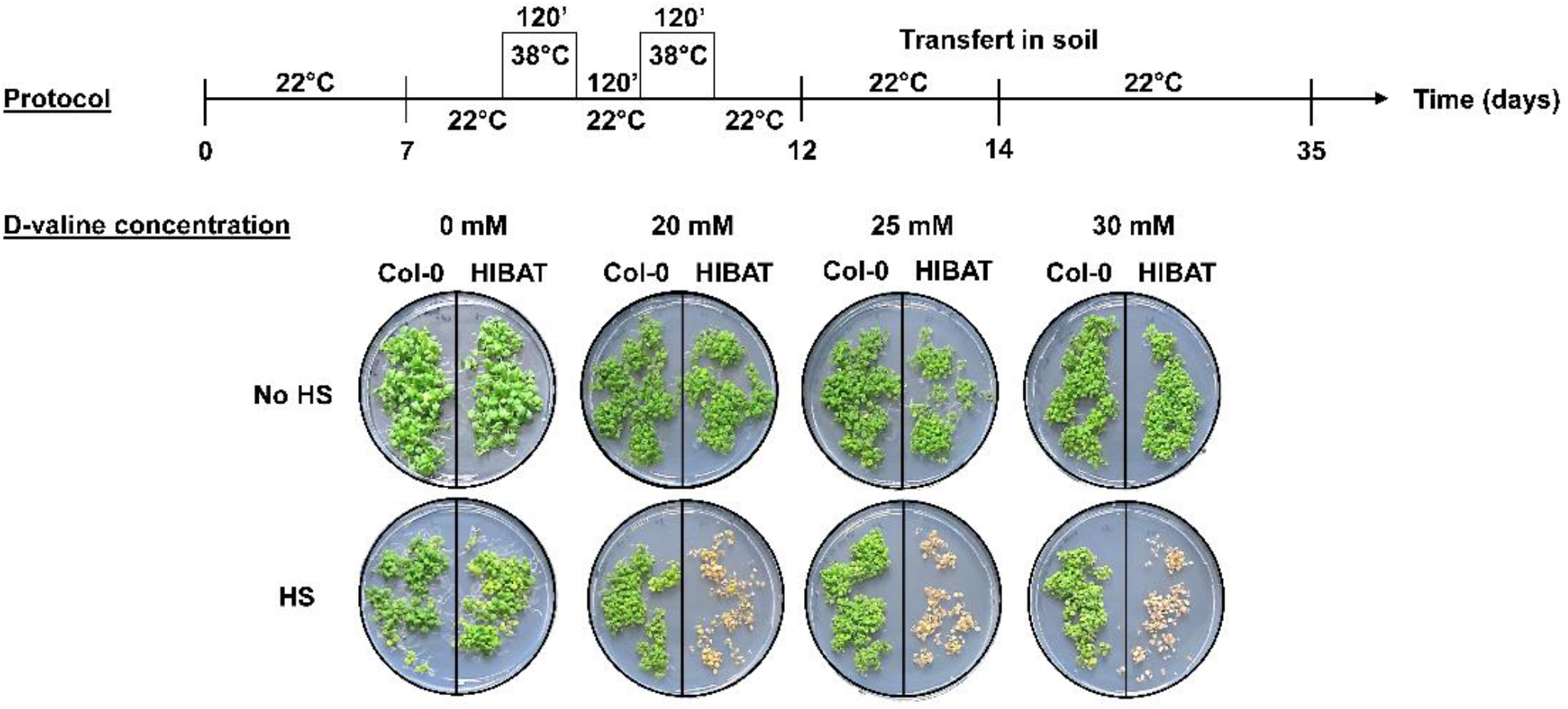
Two treatments at 38°C in the presence of D-valine kill HIBAT but not Col-0 seedlings. **A**) protocol of two weeks-old Col-0 and HIBAT seedlings grown at 22°C in the presence of increasing D-valine concentrations, then treated by two consecutive 120 min heat-treatments at 38°C, interspaced by 120 min at 22°C. **B**) Picture of Col-0 and HIBAT seedlings lawns grown on Petri dishes in the presence of 0-, 20-, 25-, and 30-mM D-valine, without (no HS) or following the two heat-treatments (HS). HIBAT seedling death was observed 6 days after the heat-treatments.

### nLUC-DAO expression is heat-specific

To screen for mutants affected in heat-sensing and heat-signaling, it is essential that the recombinant conditional HSP17.3B promoter will be specific for heat and remain unresponsive to other stressors. 12-day-old HIBAT seedlings were exposed for 5 hours, either to 250 µM H_2_O_2_, 150 mM NaCl, 300 mM mannitol, or 1 mM FLG22, at various indicated temperatures and nanoluciferase expression levels were measured in seedling’s extracts (Fig 3). In general, other stressors did not induce nLUC at 22°C and mildly affected the maximal heat-induced accumulation at 35 and/or 38°C: externally applied H_2_O_2_ showed a very minor activatory effect at low temperature, which was, however, a thousand times lower than the maximal level of nLUC-DAO, that accumulated following 2 hours at 36°C. H_2_O_2_ mildly affected heat-induced nLUC-DAO expression at 38°C (Fig 3A). Osmotic stress with mannitol did not have a significant effect at 22°C, but significantly reduced the heat-induced accumulation of nLUC-DAO (Fig 3B). NaCl stress had no activating effect at low temperatures, but reduced nLUC-DAO levels at 35°C (Fig 3C). Additionally, flagellin FLG22 did not cause measurable effects at 22°C, but reduced heat-induced nLUC-DAO accumulation at 35°C (Fig 3D). Cooling and chilling at 0°C for 5 hours had no influence on nLUC expression (Fig 4). Likewise, external application of varying concentrations of Indole-3-acetic acid, Jasmonate, Methyl-Jasmonate, Epibrassinolide, and Salicylic acid did not affect transgene expression at 31°C. Only Abscisic acid had a minimal activation effect (Fig 5 A, B). The HSP90 inhibitor radicicol (50 µM) had no effect at 22°C, and a minor yet significant co-activatory effect at 31°C (Fig 5 A). This is consistent with reports in *Arabidopsis* (Yamada et al. 2007), animal, yeast and moss cells, in which the specific binding of HSP90 inhibitors, such as radicicol, was shown to cause the dissociation from HSP90s of bound clients, such as the inactive HSF1 the cytosol (Neckers and Workman 2012). However, a mild isothermal HSR activation by radicicol and 22°C was not observed here. Even a 2-hour preincubation with 50 µM radicicol, neither enhanced at 22°C, nor significantly reduce the plant’s ability to respond to a subsequent heat shock at 37°C. The accumulation of heat-induced nLUC-DAO was nearly identical with or without radicicol preincubation (Fig 5A). This suggests that the mere radicicol-induced presumed dissociation of bound HSP90s from inactive HSF1, does not suffice to drive the activation of a heat-shock like isothermal response, as generally though (Hentze et al. 2016).

**Figure 3:**
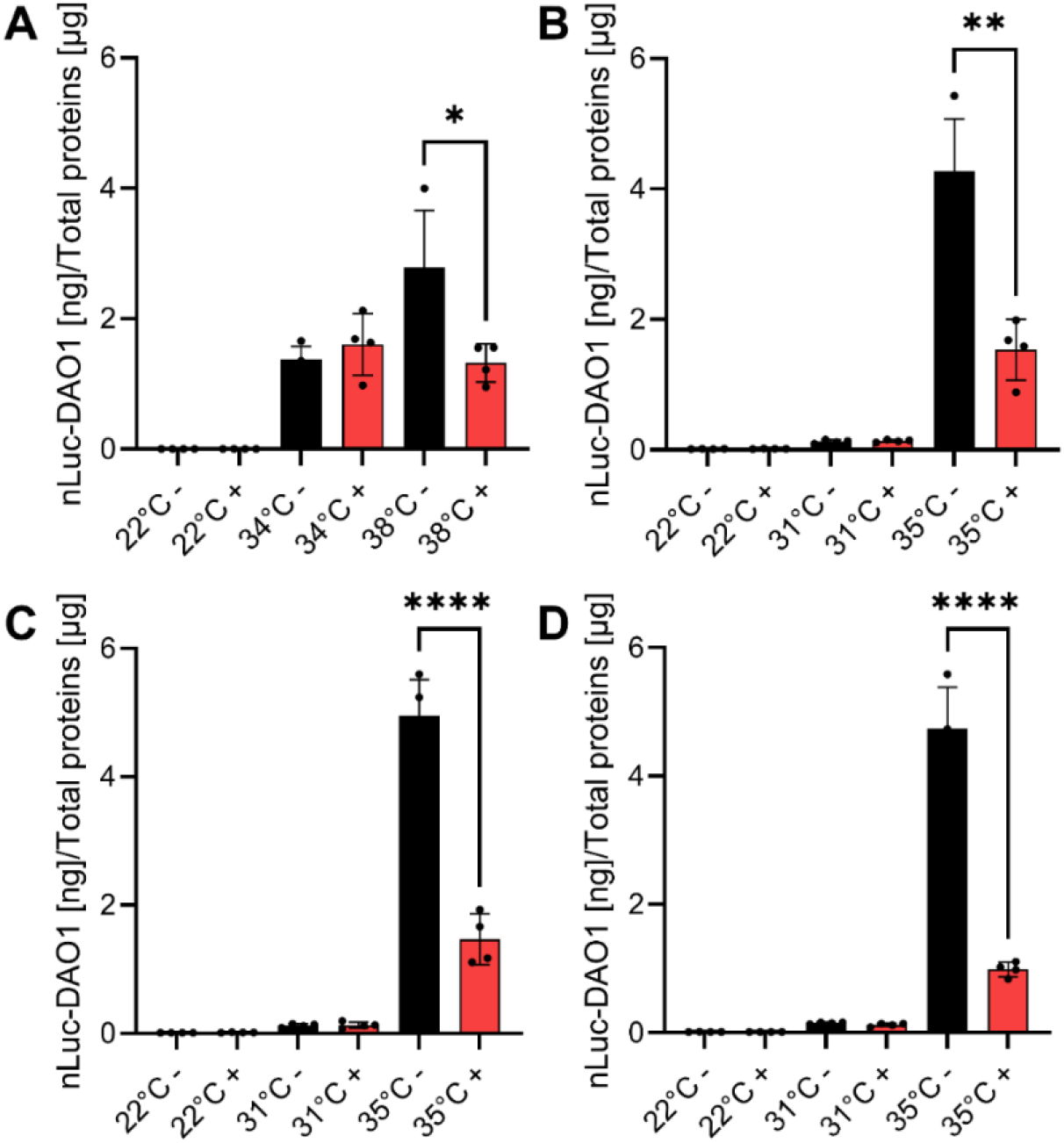
Effect of chemicals on of nLUC-DAO1 expression at different temperatures. **A**) H2O2. **B**) Mannitol. **C**) NaCl. **D**) Flagellin. 22 Seedlings on day 12 were exposed either to 250 µM H2O2, 300 mM mannitol, 150 mM NaCl or 1 mM of FLG22, for 5 hours at 22°, followed by 2 hours at the indicated temperatures, then 2 hours at 22°C. Asterisks indicate statistically significant differences determined by student T.test (*, P<0.05, NS, No Significant). Means ± S.D. (n = 4). Black: absence of H2O2, Mannitol, NaCl and FLG22. Red: presence of H2O2, Mannitol, NaCl and FLG22. Asterisks indicate statistically significant differences determined by Student t-test (* P<0.05, ** P<0.01, *** P<0.001, **** P<0.0001, NS, Not Significant).

**Figure 4:**
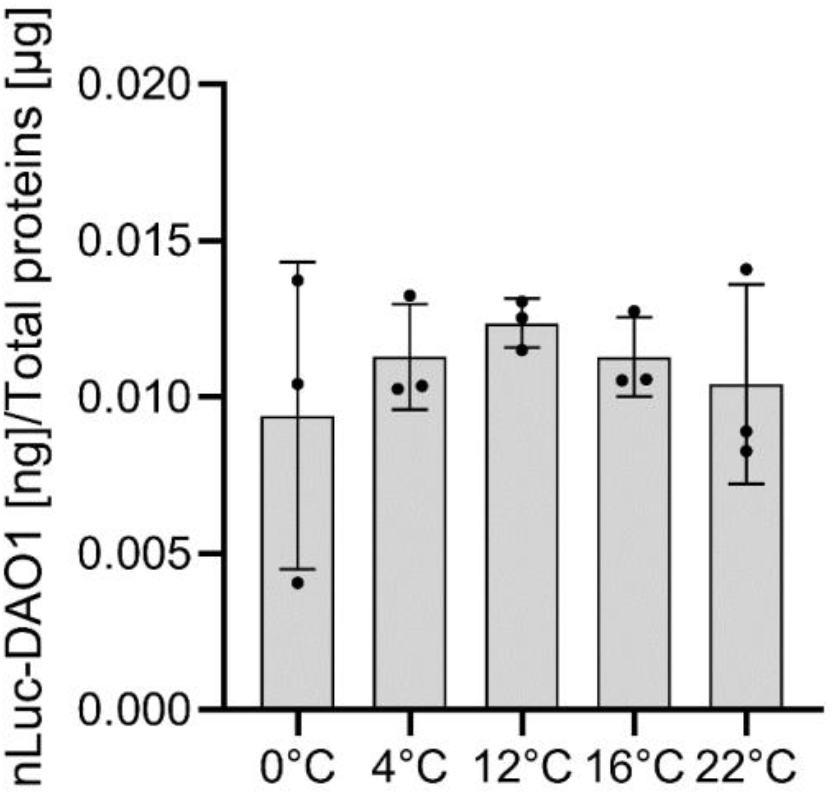
Effect of chilling and cold-shock on the expression of nLUC-DAO1 in HIBAT. Seedlings on day 12 were exposed for 2 hours to cold shock (0°C), chilling (4°C to 16°C) or at room temperature (22°C). Measures were taken from cell extracts following 2 hours of post-treatment at 22°C. means ± S.D. (n = 4). A lack of significant difference was observed at 22°C compared to all other conditions and determined by the student t-test.

**Figure 5:**
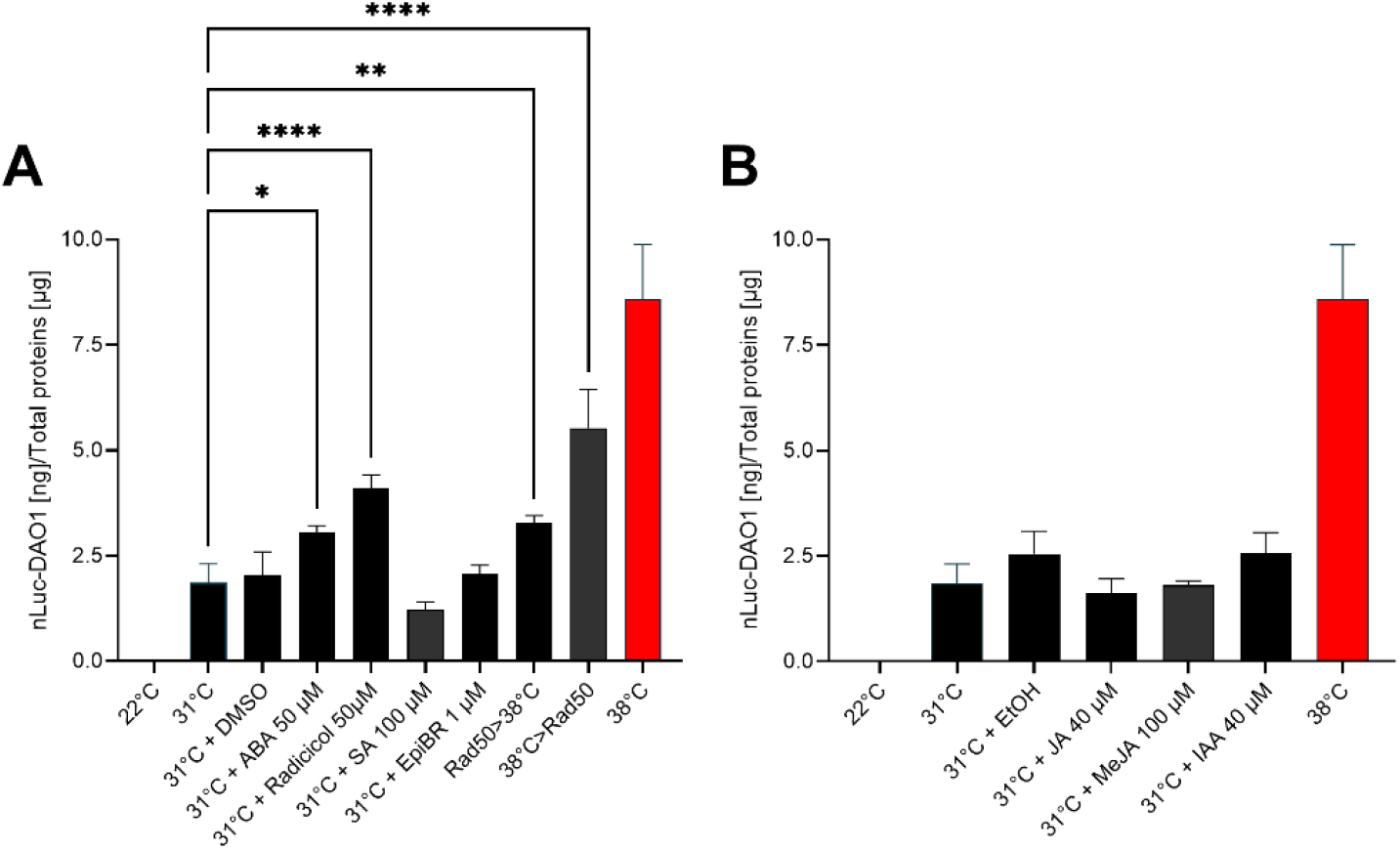
Effect of phytohormones on the expression of nLUC-DAO. Seedlings were exposed on day 12 to a 2- hours heat shock at 38°C,or to different phytohormones during 4 hours at 31°C or without any treatment at 22°C. **A**) Differential expression of the transgene in the presence of DMSO (mock), 50 µM ABA, 50 µM Radicicol, 100 µM salicylic acid (SA), 1 µM Epibrassinolide (EpiBR), 50 µM Radicicol at 22°C for 2hours then HS at 38° or 38°C Heat Shock for 2 hours and addition of 50 µM Radicicol. **B**) Differential expression of the transgene in the presence of ethanol (EtOH, mock), 40 µM Jasmonic Acid (JA), 100 µM Methyl-Jasmonate (JA) or 40 µM of Indole-3-acetic acid (IAA, auxin). Asterisks indicate statistically significant differences determined by ordinary 1-way ANOVA (* P<0.05, ** P<0.01, *** P<0.001, **** P<0.0001, NS, Not Significant).

Interestingly, the transgene was inserted 834 nucleotides downstream to the AT1G69600 gene, encoding for ZFHD1, which is a member of the zinc finger homeodomain transcription factor family. ZFHD1 is induced by drought, high salinity, and abscisic acid upon binding to the promoter region of the early responsive to dehydration stress gene ERD1 (Tran et al. 2004). RNAseq analysis (see below) showed that AT1G69600 low basal expression level is also induced about twice by heat, whereas the transgene expression was strictly depended on heat and not on drought, high salinity, or abscisic acid.

### Profile of HIBAT gene expression following heat or D-valine treatments

We next investigated whether HIBAT displayed similar a profile of heat-induced mRNA expression as the wild-type Col-0 parental strain. RNA-Seq analysis of HIBAT seedlings grown at 22°C, without or following 40-minute at 38°C, produced about 25’000 identified transcripts, of which 2’668 were significantly upregulated. Together, they contributed to a net mRNA gain of 23%, corresponding to 230’000 Transcript Per Million (TPMs). 2’159 transcripts became also significantly degraded during the short heat treatment, which together contributed to a net loss of ∼28’000 TPMs. Noticeably, many of the most heat-degraded transcripts encoded for chloroplast imported proteins. Yet, because their relative degraded amounts greatly varied between samples, the statistical analysis excluded a total of ∼200K TPMs with excessive p-values from the list heat-degraded TMPs.

Of the 18 detected genes encoding for HSP20s, which are alpha-crystalline-domain containing chaperones (Finn et al. 2014), 12 were within the 100 most extensively heat-induced genes of the entire *Arabidopsis* genome (Supplementary Data S1). Interestingly, these 12 HSP20 genes were virtually unexpressed at 22°C. Other highly upregulated genes in response to heat also included several members of the HSP90, HSP100, HSP70, and HSP60 chaperone families, as well as their co-chaperones (collectively referred to as the chaperome) (Fig 6).

**Figure 6:**
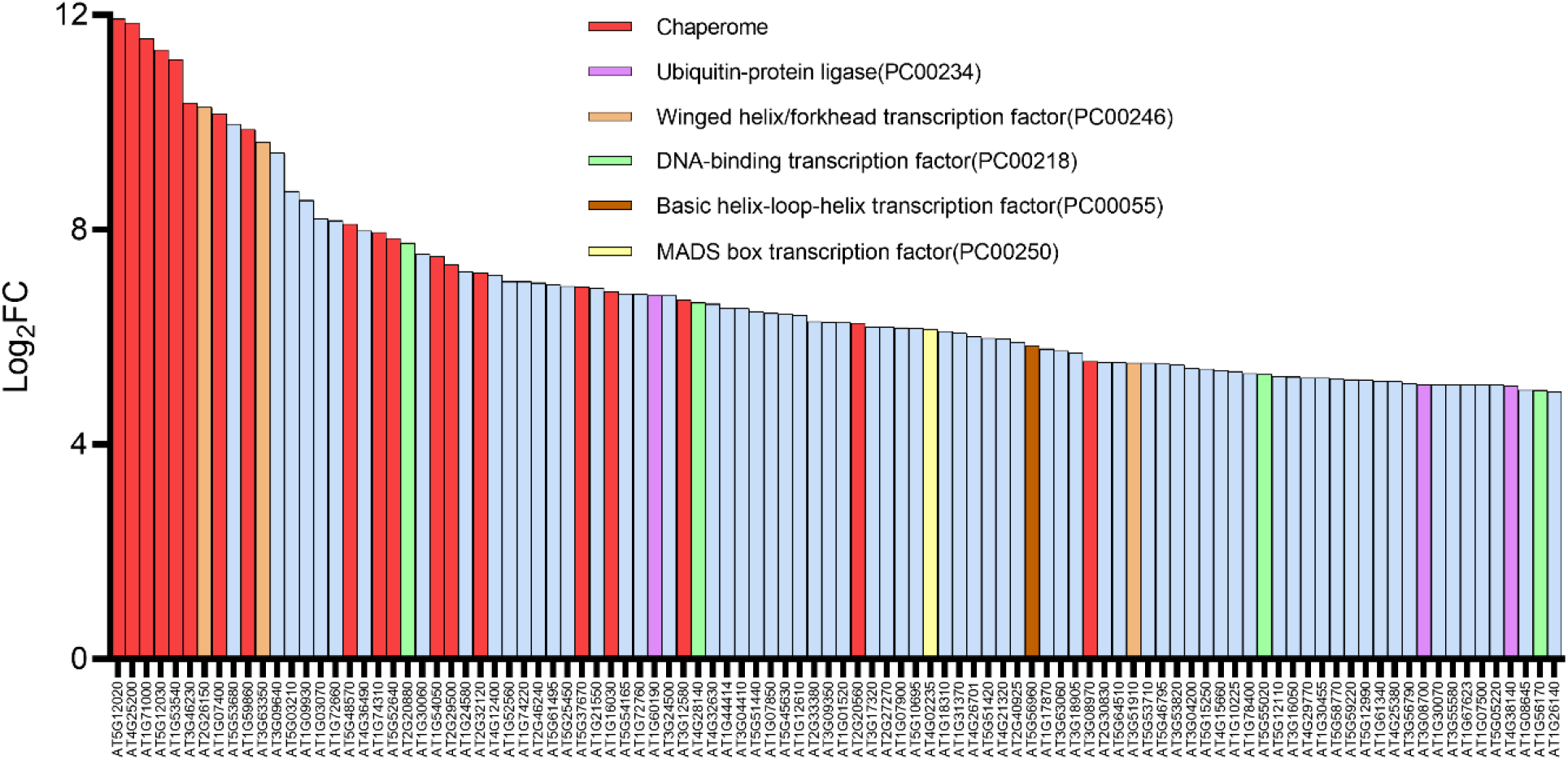
Top 100 heat-stressed genes in Arabidopsis thaliana HIBAT seedlings. The 100 most heat induced genes expressed in Log2FC (Fold-Change) corresponding to number of Transcripts per million (TPM) following 40 min at 38°C, divided by the number of TPM in untreated 14-day-old HIBAT A. thaliana seedlings at 22°C. The columns include genes encoding for chaperones and cochaperones (chaperome) (red), Ubiquitin ligases (purple), HSFs (orange), DNA-binding transcription factors (green), helix-loop-helix transcription factors (brown), MADS-box transcription factors (yellow), and others (blue). Genes families were assigned by PANTHER gene analysis (Thomas et al. 2003).

Compared to the 40 min heat-treatment that significantly accumulated the transcripts of ∼3’000 genes, and significantly degraded the transcripts of ∼2’600 genes, the presence of 25 mM D-valine at 22°C had only a minimal effect on the plant mRNA expression profile: it caused a significant, yet very mild accumulation of merely 31 transcripts, and significantly reduced 78 transcripts (Supplementary Data S1). Importantly, none were among the 100 most heat-induced or heat-degraded transcripts. Thus, RNAseq analysis confirmed that the presence of 20 mM D-valine in the growth medium have minimal to insignificant effects on the plant physiology.

RNAseq analysis from heat-treated HIBAT showed a strong correlation with two published studies using similarly heat-treated Col-0 seedlings (Blair et al. 2019; Grinevich et al. 2019) (Fig 7). A correlation analysis between the Log_2_ Fold-change in heat accumulated versus unstressed transcripts, between the current HIBAT study and two published studies (Blair et al. 2019; Grinevich et al. 2019) showed a high positive correlation for all the analyzed genes, with a Spearman correlation coefficient (*ρ)* of 0.8, 0.85 and 0.84, respectively (Fig 7 A, B, and C). When considering only chaperome genes (in red), the correlations were 0.87 and 0.92 (Fig 7 A, B, and C in red). This implies that heat shock response expression profiles of HIBAT and wild type Col-0 are very similar.

**Figure 7:**
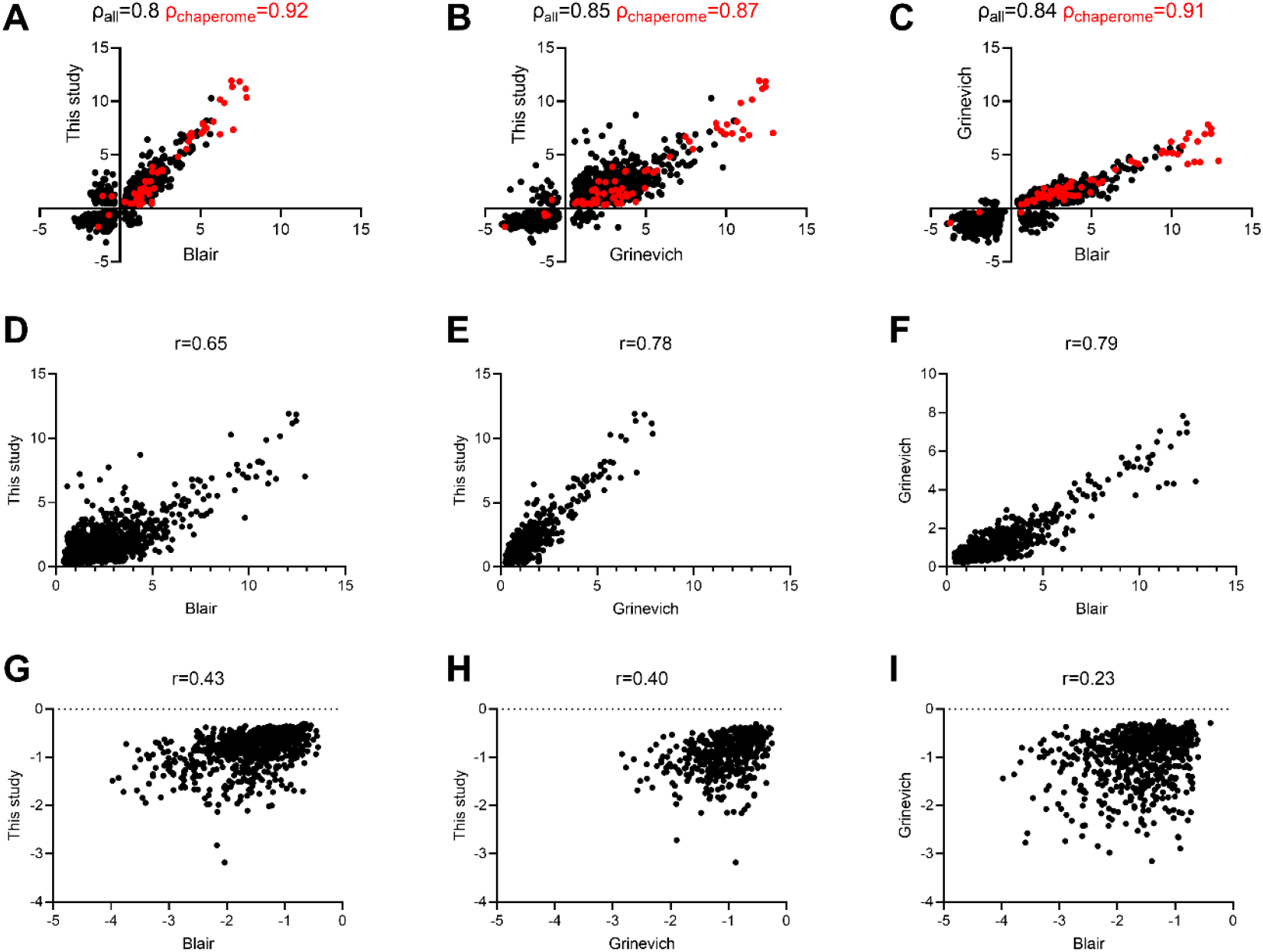
Scatter Plot showing a correlation in Log2 fold-change expression between two pairs of datasets. A): Correlation between this study and Blair et al., B) Grinevich et al., and C) between Blair and Grinevich. Spearman’s rank correlation coefficient (ρ) is indicated in black for all genes and red for the chaperomes. D-F): All upregulated genes. G-I): All downregulated genes.

#### HIBAT is an effective tool for genetic screens

##### Selection of loss-of-HSP repression mutant at low temperature

A mutation in the chromatin-remodeling complex Arp6 has been found to cause the partial dissociation of special histone H2.AZ from a recombinant plant HSP70 promoter, alongside its apparent mild de-repression at 18°C, that produced higher-than-background levels of expressed luciferase (Kumar and Wigge 2010). However, the identity of the repressors of HSP genes at basal temperatures in plants is still largely unknown. Here, the HIBAT line was used to isolate mutants impaired either in their ability to repress HSP expression at 22°C or in their ability to accumulate HSP expression following heat-treatments (Fig S6).

Following EMS mutagenesis of HIBAT seeds, a total of 3’871 M2 seedlings were screened in the absence of D-valine and in the presence of furimazine for abnormally high bioluminescence at 22°C. One mutant was identified, named “L5” that displayed significantly higher levels of nLUC-DAO both at 22°C and 31°C, compared to the parental HIBAT line. Western blots confirmed that not only the basal levels of the transgene product, but also the levels of endogenous sHSP17.7 and HSP101 proteins were significantly higher at 22°C and 31°C than in the WT HIBAT. Interestingly, at 35°C, the levels of these HSPs were mildly lower in L5 than in HIBAT (Fig 8), together suggesting that L5 is a hyper-thermosensitive mutant. We then pooled the offspring of the mutant and performed a whole-genome DNA sequencing, in an attempt to identify the causal SNPs for the observed dysregulation in the heat shock response (HSR) phenotype at 22°C and 31°C. The analysis of the 234 SNPs that were found, 16 caused stop codons or amino acid changes in exons (Supplementary Data S1). Investigation of several T-DNA insertion lines corresponding to promising SNPs, revealed one very interesting candidate, corresponding to a mutation in the HSP70-3 gene (AT3G09440), where alanine 161, which is highly conserved in all the nucleotide-binding domains of canonical eukaryotic HSP70s and bacterial DnaKs, was mutated in L5 into threonine (Fig 9). Remarkably, we observed by western blot that a T-DNA KO mutant of the HSP70-3 gene similarly expressed higher levels of HSP17.7 than in HIBAT, both at 22°C and 31°C, thereby reproducing the hyper-thermosensitive phenotype of L5 (Fig 9A).

**Figure 8:**
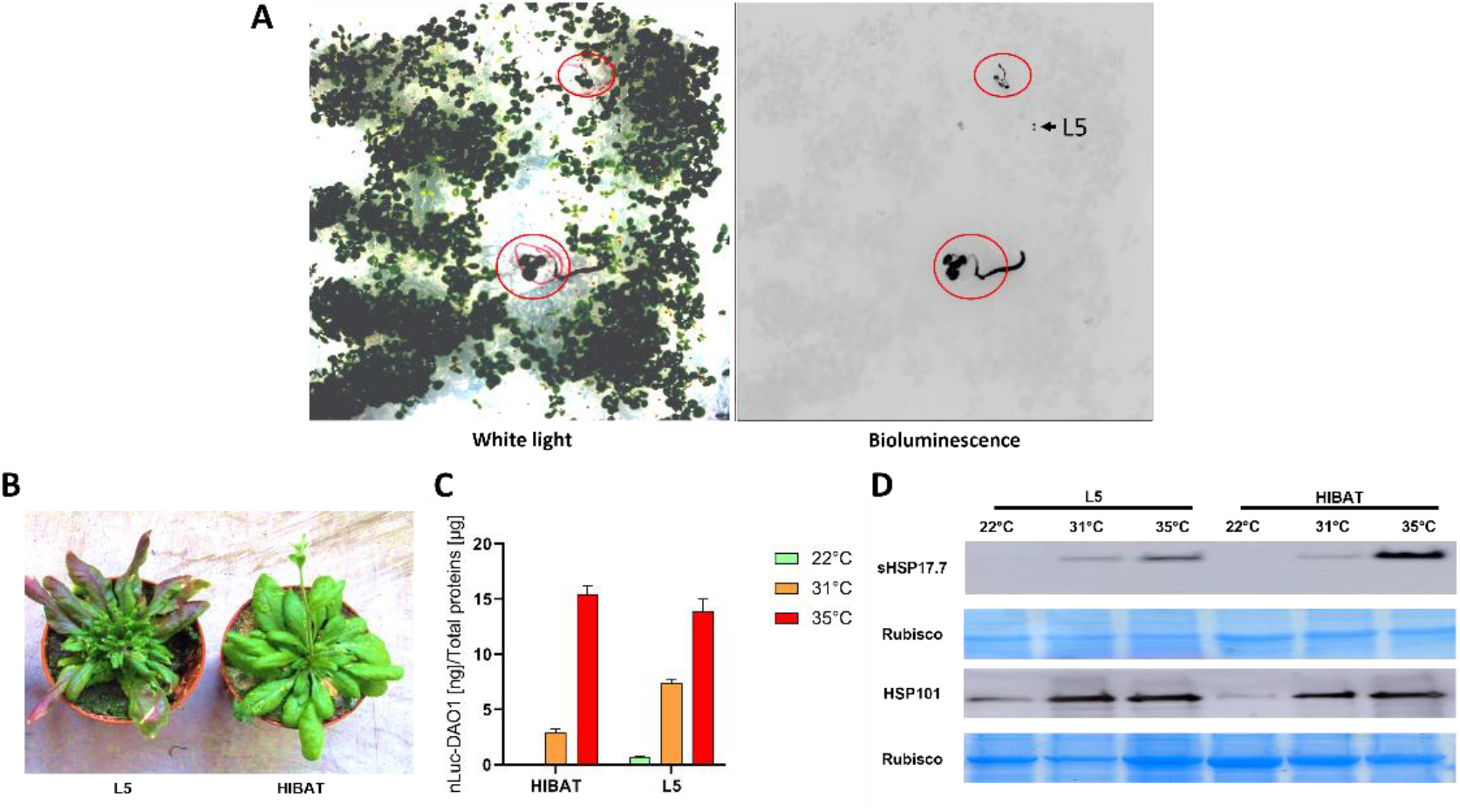
The L5 mutant is Hyper thermosensitive. A) Control pre-heat treated HIBAT seedlings (red circles) and L5 mutant (arrow). B) Phenotypes observed in the parental HIBAT line and the L5 candidate at 5 weeks old. C) 5-week-old leaves from the parental HIBAT line and the L5 candidate were exposed to 22°C, 31°C, or 35°C for 2 hours followed by 2 hours of post-recovery at 22°C. D) The expression of nLUC-DAO1, sHSP17.7, and HSP101 were then analyzed. The homogenous loading for each protein sample is represented by Coomassie blue staining of RubisCO.

**Figure 9:**
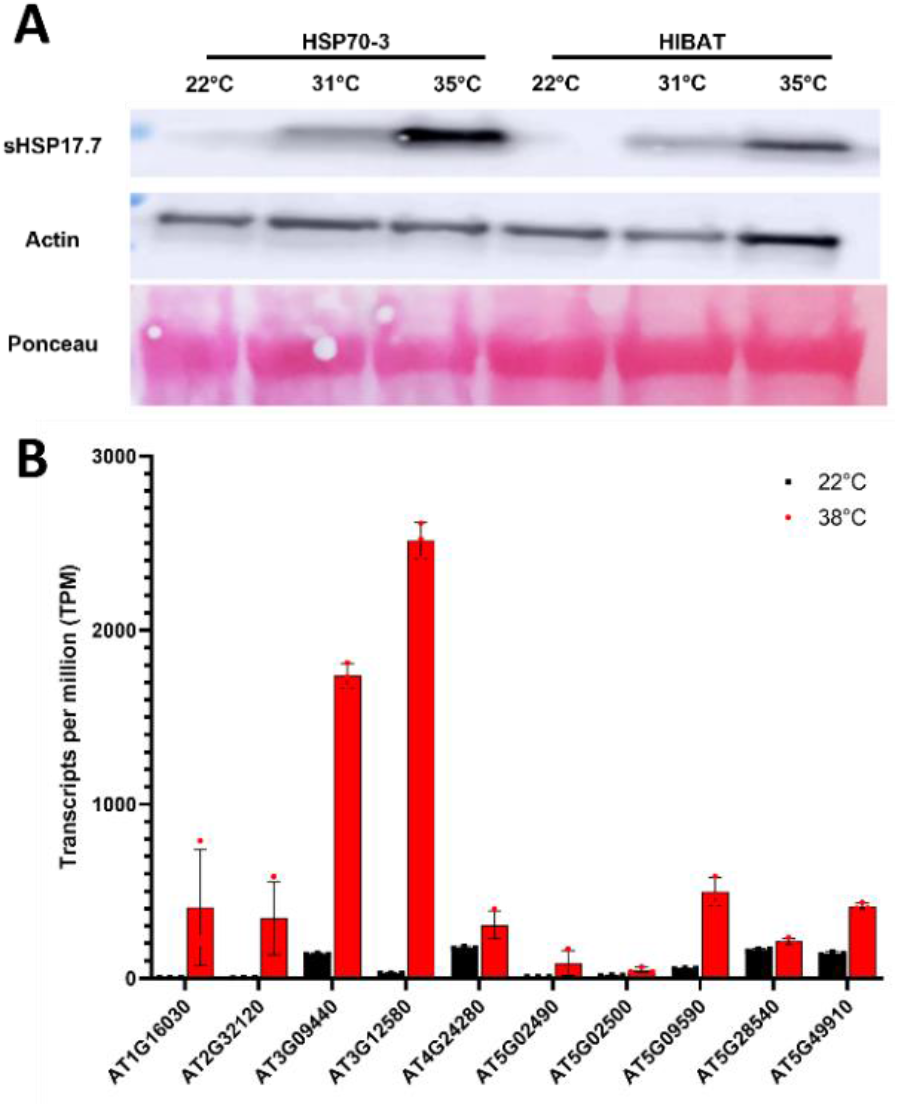
HSP70-3 expression level in response to heat stress in Arabidopsis leaves. A) Accumulation of HSP17.7 in HSP70-3 mutant line and Col-0 from 5-week-old leaves exposed to 22°C, 31°C, or 35°C for 2 hours followed by 2 hours of post-recovery at 22°C. B) Expression level of all detectable HSP70 in TPM in our transcriptomic dataset, with HSP70-3 being AT3G09440.

##### The selection of loss-of-heat-sensing/signaling putative mutants

T2 descendants of EMS-pretreated HIBAT seedlings that survived iterative heat-shocks on D-valine were further grown for 14 days in soil (without D-valine) at constant 22°C and then submitted, or not, to 40 min heat-treatment at 38°C (figure S6). nLUC activity was measured in leaf extracts and immunoblots performed with HSP17.7 and HSP101 antibodies (Fig 10). About 5% of the seedlings that survived HS on D-valine, presumably owing to their ineffective heat-induced expression of the toxic transgene, remarkably, also displayed a deficient heat-induced expression of HSP101, which is encoded by a different chromosome. The HSP17.7 antibody may recognize several closely related orthologous cytosolic HSP20s, such as AT1G07400, AT1G53540, AT3G46230. This notwithstanding, their level in the candidate mutants was also decreased, indicating that globally, cytosolic HSP20s also failed to accumulate in response to heat (Fig 10). Notably, the transgene was positioned on chromosome 1, about 6.2 million base pairs (Mbp) away from AT1G53540 and 23.9 Mbp away from AT1G07400, whose promoter and amino acid sequence are most closely related to the soybean HSP17.3B gene. Further analysis is now needed to identify in the selected putative mutants, the physiological processes, and the affected gene(s) that are responsible their loss-of-heat-sensing and signaling phenotype.

**Figure 10:**
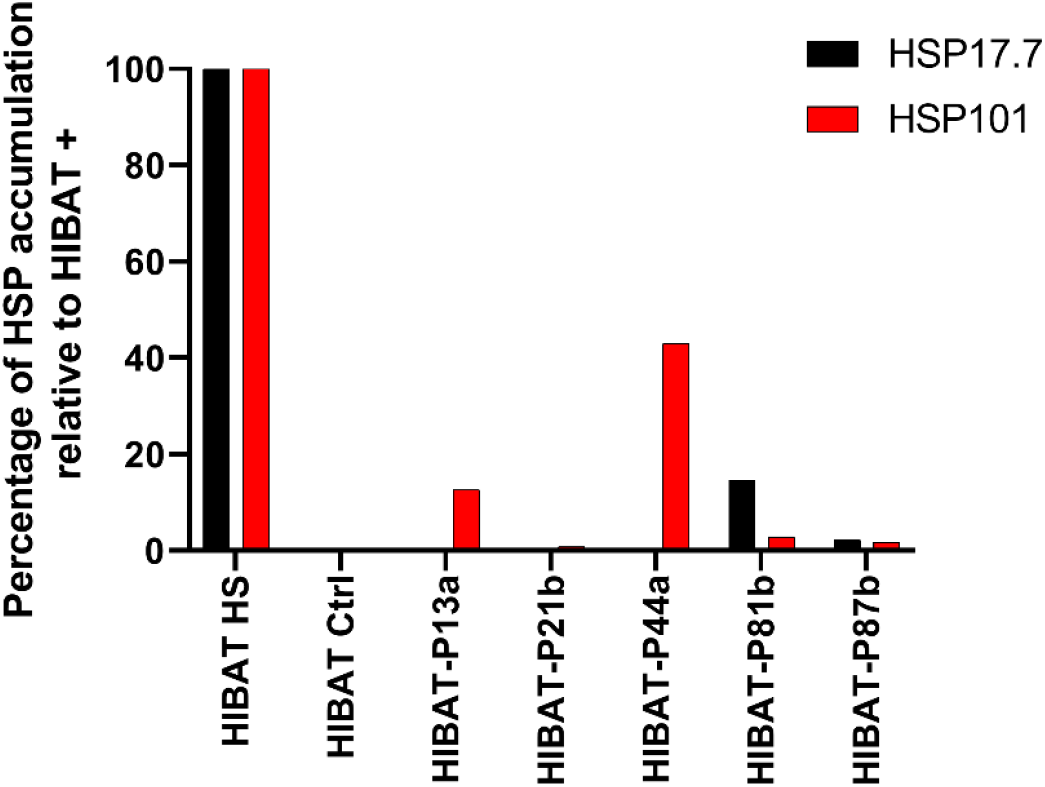
Accumulation of HSP17.7 and HSP101 in selected M3 HIBAT mutants under heat shock. Accumulation of HSPs in percentage observed by Western blot on M3 HIBAT candidates and normalized by the intensity of HSP detected in HIBAT+ line after Heat Shock. Accumulation of HSPs were normalized against HIBAT+ line and by using expression of RUBISCO in graphs. The quantification was performed by ImageJ Software (Hartig 2013).

## Discussion

### Global warming and the role of HSPs

The increasing concerns about global warming emphasize the urgent need to understand the molecular mechanisms by which higher plants, crops in particular, sense and respond to extreme temperature increments. High temperature stress can severely affect plant growth and productivity and plant’s capacity to sense upcoming heat stresses and timely buildup proper HSP-based molecular defenses, is essential for their survival (Wahid et al. 2007). The heat-shock proteins (HSPs) belong to a very heterogeneous ensemble of proteins carrying diverse, albeit complementary, stress-protective cellular functions. Whereas, by definition, all HSPs share the ability to specifically accumulate in cells, in response to a mild warming, a stronger heating and/or a severe heat-shock (Wang et al. 2004), many, although not all, of the massively heat-accumulating proteins are also molecular chaperones. They mainly belong to five highly conserved, very ancient chaperone families: the HSP60s, HSP70s, HSP90s, HSP100s and HSP20s (small HSPs), forming together with their co-chaperones, a gene category referred to as “the chaperome” (Joshi et al. 2018). Thus, given that the synthesis of two third of the plant chaperome is not induced by heat, it is misleading to globally refer to chaperome members as HSPs (Finka et al. 2011). Despite this, both the constitutively-expressed and the heat-induced chaperones can, in general, prevent the misfolding and aggregation of thermolabile proteins during heat stress, and in addition HSP60s, HSP70s, HSP90s, HSP100s that hydrolyze ATP, can “repair” already formed aggregates (Boston et al. 1996). Other heat-induced proteins, which are not molecular chaperones, such as ascorbate peroxidase, are involved in the detoxification of ROS. Others, such as HSFA2, are heat-shock transcription factors. There are also signaling molecules (calmodulin, calmodulin kinases) and enzymes producing thermo-protective and ROS-quenching metabolites (Guihur et al. 2022a; Havaux 1998). The accumulation of HSPs and metabolites is therefore key for the onset of plant survival to heat stress, and a better understanding of the molecular mechanisms underlying heat-sensing, signaling and the buildup of molecular responses, is central to develop strategies to ameliorate plant heat tolerance.

### Characterization of HIBAT, a Heat Shock Reporter Line

To investigate the molecular mechanisms leading to the accumulation of HSPs in response to elevated temperature, we generated the transgenic *Arabidopsis* reporter line HIBAT. It featured the heat-inducible HSP17.3B promoter derived from soybean (Saidi et al. 2005) driving the conditional expression of a fusion transgene composed of nano luciferase and D-amino acid oxidase (nLUC-DAO). Upon heat-induced expression of the transgene, the N-terminal domain of nanoluciferase (nLUC), HIBAT was found to generate a strong and stable bioluminescence signal. This signal correlates with the levels of endogenous heat-accumulated HSP20 proteins, while expression of the C-terminal DAO domain was found to be conditionally toxic. However, this toxicity only occurs in the presence of D-valine, which results in the production of harmful reactive oxygen species (Smirnoff and Arnaud 2019; van der Eerden 1982; Ogier de Baulny and Saudubray 2002; Taylor et al. 2004). Due to the presence of a single copy of the transgene, in a non-coding intergenic region, HIBAT did not express any detectable nLuc-DAO and consequently grew unaffected by D-Valine at 22°C. In contrast, 98% of the HIBAT seedlings did not survive heat treatments on D-Valine (Fig 2B, Fig S5). Heat-treatment without D-valine also accumulated endogenous HSP17.7 and HSP101 in heat-treated Col-0 and HIBAT seedlings, but not in quadruple *hsfa1* mutant that cannot produce HSPs in response to a heat-shock (Fig 1B) (Liu et al. 2011).

RNA-seq analysis of untreated and heat-treated HIBAT seedlings confirmed that like in the wild-type Col-0, most cytosolic members of the HSP20 family are virtually unexpressed under basal growth temperature, and dramatically overexpressed under heat-shock (Saidi et al. 2005; Bourgine and Guihur 2021; Guihur et al. 2021). Twelve out of approximately twenty HSP20 genes were among the top 100 most heat-induced genes of the plant genome. Considering that the HSP20 genes are ∼1/1500 of the total *Arabidopsis* genome, this is a 180-fold over-representation of this gene family among the 100 most heat-induced genes of the plant.

The very tight repression of the HSP20s expression that was observed at 22°C suggests that although HSP20s are essential for plant recovery from heat stress, their futile expression at low temperature would be, for unclear reasons, detrimental to unstressed plants (Guihur et al. 2022b; Wu et al. 2022; Guihur et al. 2021; Guihur et al. 2022a; Sun et al. 2016; Bourgine and Guihur 2021). In contrast, other families of heat-induced chaperones, such as the HSP60s, HSP70s, HSP40s, HSP90s, and HSP100s, which have orthologs with high levels of constitutive expression at low temperature (e.g., chloroplastic HSP70, AT4G24280, Fig 9B), are proportionally only mildly heat-induced. But this may be a misconception that is produced from a deliberate choice to express the effect of heat on the expression of a given gene, as the log of <u>fold change</u> of its heat-accumulated transcripts per million (TPMs), divided by its ground temperature TPMs, rather than as a net <u>difference</u> between the heat-accumulated TPMs and the ground temperature TPMs.

### The HSP17.3B promoter is highly specific heat

Identical heat-treatments can produce much more HSP mRNAs in dehydrated plants, than in hydrated plants (Zandalinas and Mittler 2022; Cohen et al. 2021). Yet, for an *Arabidopsis* reporter line designed to screen for mutants affected in plant heat-sensing and signaling, we sought for an expression system that is specific to heat with minimal interferences from other stressors. Indeed, the synthesis of the reporter protein in HIBAT was found to be specific to heat with no significant interferences observed for salt stress (NaCl) and osmotic stress (mannitol), although both are thought to mediate Ca^2+^- and H_2_O_2_-mediated cytosolic signals that also lead to HSPs accumulation (Belver and Travis 1990; Huang et al. 2016). Pathogen-like stress (flagellin-22), and various phytohormones remained ineffective, except for a minor activatory effect by abscisic acid, confirming previous evidence that abscisic acid is necessary for the onset of plant thermotolerance (Suzuki et al. 2016; Huang et al. 2016). A similar very minor activatory effect was found for H_2_O_2_ which participates in the activation of HSFs and acts in various organisms as a secondary messenger, leading to HSP expression (Volkov et al. 2006; Davletova et al. 2005; Maruta et al. 2012; Černý et al. 2018). We did not observe an effect of cooling and chilling (Guo et al. 2018), but this is not unexpected from more rigid plasma membranes, expected to freeze up more than activate the CNGC2/4 heat sensors.

### The effect of radicicol: implications for HSF1 activation

Concurring previous reports in mosses and other organisms (Neckers and Workman 2012; Hentze et al. 2016), the minor accumulation of nLUC-DAO at 31°C was co-activated by the HSP90 inhibitor radicicol (Fig 5 A). Yet, remarkably, this did not yield a maximal heat-shock response. This is suggesting that the heat-induced activation of hypophosphorylated HSF1 requires more than its mere dissociation from inhibitory HSP90s, as generally believed (Hentze et al. 2016). Indeed, experimental evidence have shown that HSF1 activation also requires a specific Ca^2+^ entry dependent signal originating from the CNGC2/4 channels in the plasma membrane. This heat-shock signal contributes to HSF1 hyperphosphorylation and the release of inhibitory chaperones, such as HSP90 and HSP70 (Saidi et al. 2011; Finka et al. 2012).

The transgene was found to be inserted close to AT1G69600, which encodes for ZFHD1. This member of the zinc finger homeodomain transcription factor family accumulates under drought, high salinity, and abscisic acid (ABA). Whereas the transgene expression was unresponsive to these stresses, the very low basal expression levels of ZFHD1, were nearly doubled in response to heat, suggesting a possible overlap between the two stresses.

The RNAseq analysis confirmed the primary importance of HSP20s in the plant response to heat stress, in line with previous observations (Guihur et al. 2021). Yet, aside from *in vitro* indications that HSP20s may generally prevent protein aggregation and possibly protect membranes from hyperfluidization (Torok et al. 2001), it is not clear what is the specific mechanism by which the heat accumulated HSP20s effectively establishes plant acquired thermotolerance. Moreover, it is not clear why at low temperature, their expression is being repressed so tightly, suggesting that HSP20s may carry specific cellular functions other than mere prevention of protein aggregation. Given that plant acquired thermotolerance also results from a prior, mildly heat-induced repression of a proapoptotic signal, which would otherwise be deadly to the heat-shocked plants (Finka et al. 2012), it is tempting to speculate that heat-pre-accumulated HSP20s may contribute to plant acquired thermotolerance by repressing otherwise deadly apoptotic signals, during a noxious heat-stress, which, if they were to be constitutively expressed in unstressed plants, would in the long term affect cellular differentiation and tissue development.

It is worth noting many chaperome genes in the plant genome were either undetected, or did not become significantly heat-accumulates, or were heat-degraded, as observed in the case of AT3G04980, AT2G41000, AT3G12170, AT5G12430, AT5G06910, and AT2G33210. This confirms that chaperones and co-chaperones should not be misleadingly referred to as heat-shock proteins (Finka et al. 2011; Guihur et al. 2021).

### Utilizing HIBAT to identify an HSR derepressed mutant

The identity of the repressors of plant HSP genes expression at basal, non-heat shock temperatures remains unclear. An *Arabidopsis thaliana* mutant in Arp6, which is a component of a chromatin remodeling complex, displayed a weaker repression by H2A.Z histones of the transcription of a cytosolic HSP70 at 18°C (Cortijo et al. 2017; Kumar and Wigge 2010). In the present study, ethyl methane sulfonate-mutagenized HIBAT seeds were cultivated at a constant 22°C (in the absence of D-valine) and treated with furimazine, which led to the identification of mutant L5, that exhibited an abnormally derepressed expression at 22°C of nanoluciferase, as well as endogenous HSP17.7 and HSP101. Compared the wild-type HIBAT parental line, L5 displayed increased bioluminescence both at 22°C and 31°C. Whole-genome sequencing of L5 revealed several potential single nucleotide polymorphisms (SNPs), including a promising SNP in the HSP70-3 gene (AT3G09440) that changed a conserved Alanine (Ala) into Threonine (Thr) at position 161 (Supplementary Data S1). This Ala residue is situated in the nucleotide-binding domain of the HSP70 protein at the center of a conserved alpha-helix. It faces R513, which is in the hinge between the protein binding base and the open C-terminal lid that in the ATP-bound state of HSP70, become tightly closed onto a bound protein substrate. Ala161 is present in all canonical HSP70 family members (DnaK-homologues) across bacteria, archaea, and eukaryotes, but varies in non-canonical HSP70- related gene families, such as bacterial HSCA/Cs and HSP110s. Two computational tools, PredictSNP (Bendl et al. 2014) and PPVED (Gou et al. 2022), both predicted a deleterious effect of an A to T substitution on protein function (Fig S7). Further, severe deleterious effects resulting from an A to T substitution within an alpha-helix of an *Arabidopsis* kinase have been reported (Piovesana et al. 2023). Whereas more research is needed to ascertain that this mutation is indeed responsible for the observed hyper thermosensitive phenotype of L5, it is worth mentioning that a T-DNA knockout *Arabidopsis* mutant of the HSP70-3 gene has been found to be defective at activating the plasma membrane phospholipase Dδ (Song et al. 2021). A detailed lipidomic analysis revealed that at low temperatures, membranes from the HSP70-3 mutant were enriched with unsaturated lipids, compared to the membranes of the parental strain (Song et al. 2021). Given that CNGC2/4 heat-sensor channels are being activated by a heat-induced increases of the plasma membrane fluidity (Finka et al. 2012; Saidi et al. 2011), a constitutively more fluid plasma membrane in the HSP70-3 mutant could render its CNGC2/4 channels more responsive to lower temperatures, leading to the production of some HSPs at 22°C and 31°C, as observed in L5. Moreover, it is tempting to speculate that if HSP70-3, which is the second most heat-inducible cytosolic HSP70 (Fig 8B), would play a central role in the repression of HSF1 activity in the cytosol at 22°C (Morimoto 1998), both L5 and the Hsp70-3 KO mutant would be expected to abnormally express some higher amounts of HSPs at lower temperature, as shown in Fig 9. More research is needed to clarify on the role of HSP70s in general, and of HSP70-3 in particular, in the repression mechanism the plant at low temperature and in the activation the heat-shock response at high temperature.

## Conclusion

This study provides novel insights into the molecular mechanisms by which land plants sense and appropriately respond to increments in ambient temperature. Our findings lay the foundation for future research using loss-of-function mutants involved in the heat-signaling, the regulation of HSP-expression, HSP chaperones in particular, and their role in the onset of acquired thermotolerance in plants. The HIBAT reporter line is a powerful and valuable tool to identify new targets involved in plant responses to heat stresses and ultimately contributes to the research aiming to ameliorate plant thermotolerance. This knowledge is essential to the design of future strategies to mitigate the dramatic adverse effects of global warming on crop plants and warrant human food security (Bita and Gerats 2013).

## Material and methods

### Generation of HIBAT

#### Transgene design and transformation of *A. thaliana* plants

The pGmHsp17.3b, *nLUC*, and *DAO1* DNA sequences (Saidi et al. 2005; Boute et al. 2016; Erikson et al. 2004) were synthesized by GenScript Biotech (https://www.genscript.com) and cloned into the destination vector pFR7m24GW (provided by Prof. Niko Geldner, University of Lausanne) containing the FastRed cassette for transgenic seed selection (Shimada et al. 2010) and was transformed into *A. thaliana* Col-0 background via *Agrobacterium tumefaciens* (strain GV3101) by floral-dip method (Clough and Bent 1998; Zhang et al. 2006).

##### Selection of transformant seeds and determination of copy number of HIBAT

The choice for the HIBAT line was based the presence of a single copy of the transgene, as first evidenced by a mendelian segregation in the T2 generation of 1:4 ratio for red transformed seeds over black non-transformed seeds. This was confirmed by whole genome sequencing (Robinson et al. 2011). Reads were blasted against to the Col-0 *A. thaliana* genome (Altschul et al. 1990) and the single T-DNA insertion was localized at position 26180497 of chromosome 1. Transgene positioning and orientation was confirmed by PCR.

##### Plant materials and growth conditions

Unless otherwise stated all seeds were in the Columbia-0 (Col-0) background. The T-DNA insertion mutant (HSP70-3, SALK_148168) were obtained from the Nottingham Arabidopsis Stock Centre (http://arabidopsis.info/). The quadruple HSFA1 KO mutant was provided by Charng YY lab (Liu et al. 2011). *A. thaliana* seeds were surface sterilized with 70% ethanol and sowed on ½ MS plate (Duchefa Biochimie, ref P14881.01), and 0.8% (m/v) plant agar (pH 5.8). Following stratification (48 h, 4°C, in the dark), plates were transferred to a continuous light growth room (22°C, 100–120 µmol m−2 s−1, 60% humidity) or long days condition (16h light/8h dark). After 2 weeks, plants were transferred to soil if necessary.

##### Physiological assays

HIBAT and Col-0 plants were grown in continuous days conditions at 22°C. Root length was measured using ImageJ (Hartig 2013) software (version 1.53) at 5 and 10 days for a total amount of 80 seedlings. Regarding fresh weight, shoots from 80 seedlings were dissociated from roots and weighted at 9, 13 or 16 days old. Finally, the percentage of germination of the seed in HIBAT and Col-0 lines were determined at 5 and 10 days old with a total amount of 100 seedlings.

##### Heat treatments

Plant seedlings were heated in a growth cabinet chamber (phytotron), incubator (microbial, for and Fig S4 and Fig 2) or a thermoblock (for Fig 1C). Respective time, intensity of temperature and age of plant used are indicated in figures (See also experimental design Fig S6).

##### Acquired thermotolerance assays

HIBAT and Col-0 *A. thaliana* lines were grown on ½ MS medium 1.2 % Agar for 14 days in continuous days conditions and plates were imaged before heat treatment. Plants were submitted to heat priming at 36°C for 2 hours followed by 2 hours of recovery at 22°C and then exposed to a noxious heat shock at 45°C for 45 min. Unprimed plants were exposed to the same noxious HS. Plants were then grown for 7 days at 22°C and plates were imaged.

##### Iterative heat stresses on D-valine

HIBAT and Col-0 plants were grown on 0 mM, 20 mM, 25 mM, and 30mM D-valine Agar plates containing ½ MS medium in a growth cabinet chamber under long day condition (16 hours light, 8 hours dark) at 22°C. At day 7, a heat program was set up to apply each day for 5 days, two consecutive 2 hours heat-shocks at 38°C interspaced by 2 hours at 22°C. At day 12, plants were allowed to recover for 2 days at 22°C and were imaged.

### Other Stress treatments

#### H_2_O_2_, Mannitol, NaCl and FLG22

2 weeks-old seedlings were transferred in an Eppendorf tube of 1.5 mL containing 250 μM H_2_O_2_, 300 mM mannitol, 150 mM of NaCl, or 1 mM of Flagellin 22 (provided by Prof. Philippe Raymond, University of Lausanne) in a final volume of 1 mL of water for 5 hours (Réthoré et al. 2019). Seedlings were then exposed for 1 hour to the different stressors at the indicated temperatures and following 2 additional hours at 22°C, nLUC activity was measured in seedling’s total proteins extracts.

#### Cold stress and chilling

2 weeks-old seedlings were transferred to .5 mL Eppendorf tube of 1 ml containing water. Cold stress and chilling were applied for 2 hours using a thermoblock (4°C, 12°C, 16°C, and 22°C) or ice (0°C), and following 2 additional hours at 22°C, nLUC activity was measured in seedling’s total proteins extracts.

##### Hormonal and radicicol treatments

Abscisic acid (ABA), salicylic acid (SA), Epibrassinolide (EpiBR) and radicicol were stored in various stock solution in DMSO, while jasmonic acid (JA), methyl-jasmonate (JA) and Indole-3-acetic acid (IAA, auxin) were stored in ethanol. 2 weeks-old seedlings were exposed for 1 hour at 22°C followed by 4 hours at 31°C followed by 2 hours of post-incubation at 22°C, except for the specific radicicol experiment where seedlings were exposed to 38°C.

##### Isolation of soluble protein and Western-Blot Analysis

Plants materials were grinded at RT by a plastic pestle in an Eppendorf tube of 1.5mL and resuspended with a BAP buffer (50 mM Tris-HCL (pH 7.5), 100 mM NaCl, 250 mM mannitol, 5 mM EDTA, 10% (v/v) glycerol, and protease inhibitor cocktail diluted at 1:300 (v/v) (Sigma-Aldrich, ref P9599). Plants extracts were centrifuged for 10 min at 12’000g at 4°C. Supernatants containing soluble proteins fraction were transferred in a new Eppendorf tube of 1.5mL and the total protein concentration was determined by BRADFORD assay (Sigma-Aldrich, ref 23238).

##### Nanoluciferase activity

nLUC-DAO1 bioluminescence was detected by using the Nano-Glo Luciferase Assay System Kit from Promega (ref 1110) and the HIDEX plate reader (version 5067). The bioluminescence was analyzed for 1-second giving count per second. Using a standard curved with purified nanoluciferase, the bioluminescence emission of total protein extracts was then converted in ng of nLUC-DAO1 per ug total protein in the extract.

#### In vivo observation of nanoLuc activity

The substrate of the nano-luciferase diluted at 1:100 (v/v) (furimazine, Kit from Promega (ref 1110)) was spread on seedling that grew on Agar plates. Light emitting seedlings were imaged using an ImageQuant LAS 500.

##### EMS mutagenesis of HIBAT seeds and screening for mutations

∼6 000 seeds were mutagenized using 0.4 % ethyl methanesulfonate (EMS) in 100 mM phosphate buffer, pH 7.5 for 8 hours, then washed several times in 100 mM phosphate buffer, pH 7.5 as previously described (Kim et al. 2006). M1 seeds were sowed in the soil to generate and harvest the M2 generation of HIBAT mutant lines in a growth chamber with long days conditions (16 hours of light, 8 hours of dark) at 22°C. Heat-treated plants showing decreased nanoluciferase levels and non-heated plants showing higher nanoluciferase levels at 22°C were retained as candidates and their phenotype was further addressed by western blot in the M3 generation. To identify the causative mutations, putative M3 candidates were backcrossed to the parental line HIBAT. F2 plants phenotypes were determined by nanoluciferase activity and western blotting. Putative candidates were separately pooled and 5μg of purified genomic DNA was used for whole genome sequencing. For each sample a ∼100-fold coverage of the *A. thaliana* genome was obtained, and SNPs were identified using the SIMPLE approach as described by (Wachsman et al. 2017).

##### RNA sequencing

2 weeks old HIBAT seedlings were exposed to 22°C or 38°C for 40 min. RNA was extracted from plants using MACHEREY-NAGEL NucleoSpin RNA plant KIT (REF 740949.50). Three biological replicates (10-15 seedlings in each replicate) for each condition were sent for RNA sequencing with a total of 1ug of RNA with the Illumina TruSeq Stranded mRNA library- It was then sequenced on the Illumina HiSeq 2500 platform. Purity-filtered reads were adapters and quality trimmed with Cutadapt v. 1.8 (Martin 2011). Reads matching to ribosomal RNA sequences were removed with FastQ Screen v. 0.11.1 (Wingett and Andrews 2018). Remaining reads were further filtered for low complexity with Reaper v. 15-065 from the Kraken suite (Davis et al. 2013). Reads were aligned against *A. thaliana*. TAIR10.39 genome using STAR v. 2.5.3a (Dobin et al. 2013). The number of read counts per gene locus was summarized with htseq-count v. 0.9.1 (Anders et al. 2015) using *A. thaliana*. TAIR10.39 gene annotation. Quality of the RNA-seq data alignment was assessed using RSeQC v. 2.3.7 (Wang et al. 2012). Reads were also aligned to the *A. thaliana*. TAIR10.39 transcriptome using STAR v. 2.5.3a (Dobin et al. 2013) and the estimation of the isoforms abundance in Transcript Per Million (TPM) was computed using RSEM v. 1.2.31 (Li and Dewey 2011). Genes with low counts were filtered out. Differential expression was computed with DESEQ2 package (Love et al. 2014). P-values were adjusted by the Benjamini-Hochberg (BH) method, controlling for the false discovery rate (FDR). The RNAseq data set have been deposited to the Sequence Read Archive (SRA) at the National Center for Biotechnology Information (NCBI) repository with the dataset identifier PRJNA948681.

##### Figures and Statistical analyses

Figures and statistical analysis were made using GraphPad Prism version 9.5.0 for Windows, GraphPad Software, San Diego, California USA, www.graphpad.com

## Acknowledgements

We thank Dr. Alexandra Waskow, John Perrin, Gizem Demirkiran, and Tatiana Fomekong Tsoutezo Mbefo for assistance with data collection and insightful discussion.

## Fundings

This work was funded by grant 31003A_175453 from the Swiss National Science Foundation to P.G. and a Swiss National Science Foundation grant n°CRSK-3_196689 to A.G.

## Author information

Anthony Guihur and Baptiste Bourgine have contributed equally to this work.

## Contributions

AG, BG, and PG designed the experiments. AG and BB performed the experiments. AG, BB, and MER analyzed the data. AG, MER, and PG wrote the manuscript. AG, BB, and MER made the figures. All authors read and approved the final manuscript.

## Corresponding authors

Correspondence to Anthony Guihur or Pierre Goloubinoff.

## Ethics approval and consent to participate

Not applicable.

## Consent for publication

All authors affirm consent for publication.

## Competing interests

All authors declare that they have no competing interests.

---

**Figure S 1 :**
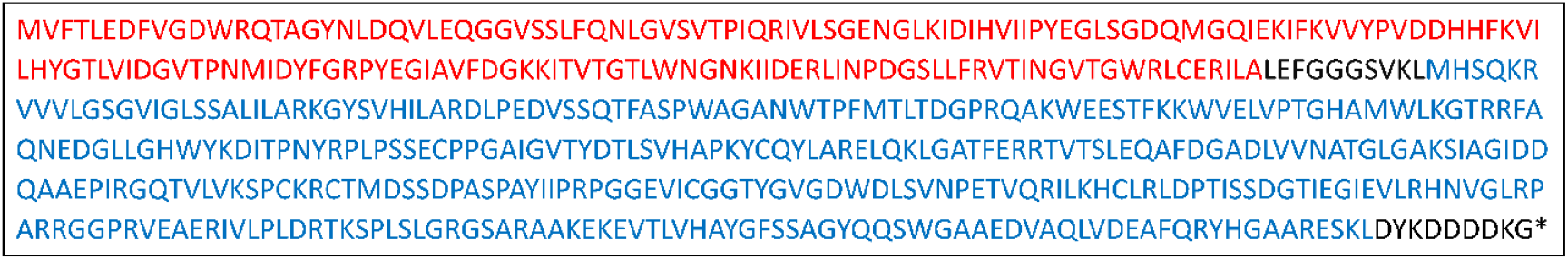
Translated protein sequence of the transgene. Red: nLUC derived from the catalytic subunit of oplophorus-luciferin 2-monooxygenase found in Oplophorus gracilirostris. Blue: DAO-1 (D-amino-acid oxidase) from Rhodosporidium toruloides. Black: Linker between DAO and nLUC and a C-terminal epitope for Flag antibodies (DYKDDDDK) (Katsuta et al. 2020). Full protein and gene sequences in (Supplementary Data S1).

**Figure S 2 :**
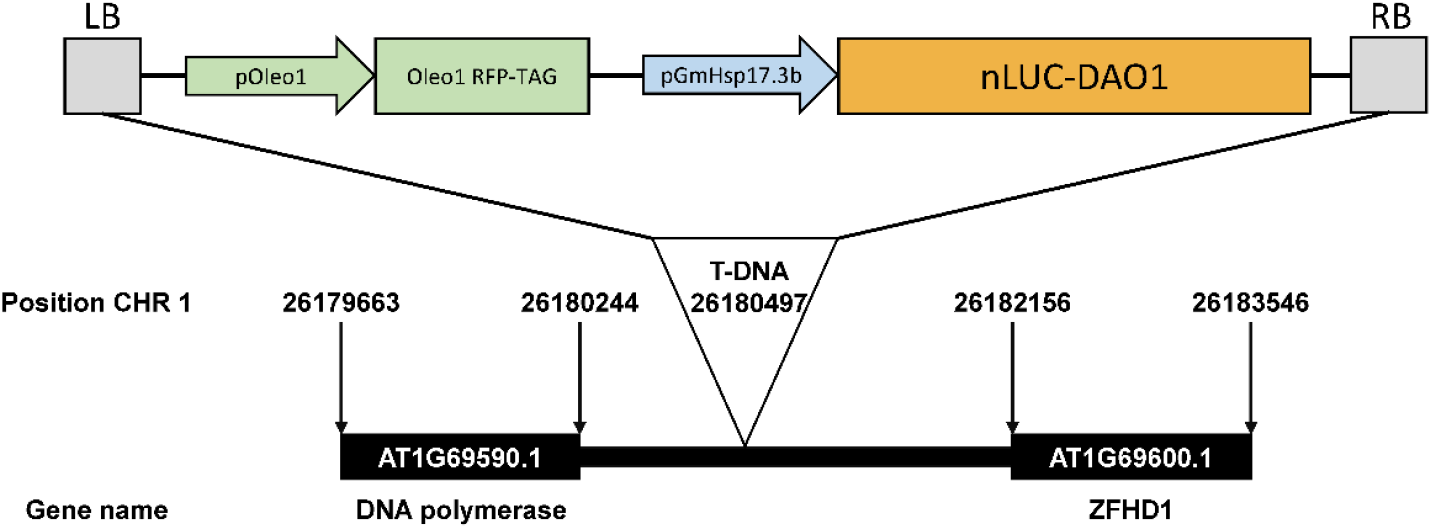
Scheme of the transgene. LB: left border, pOleo1: promoter of OLEOSIN 1 (Zhong et al. 2020), Oleo RFP-TAG: constitutive expression of a red fluorescent TAG (RFP-TAG), coupled with OLEOSIN 1 (Oleo1 RFP-TAG), pGmHSP17.3: soybean heat-inducible promoter (Treuter et al. 1993), nLUC-DAO1+Flag (DYKDDDDK epitope): nLUC is a novel and versatile small bioluminescence platform (England et al. 2016), DAO1: D-amino acid oxidase that act as a conditionally toxic negative marker in the presence of D-valine (Gisby et al. 2012).

**Figure S 3 :**
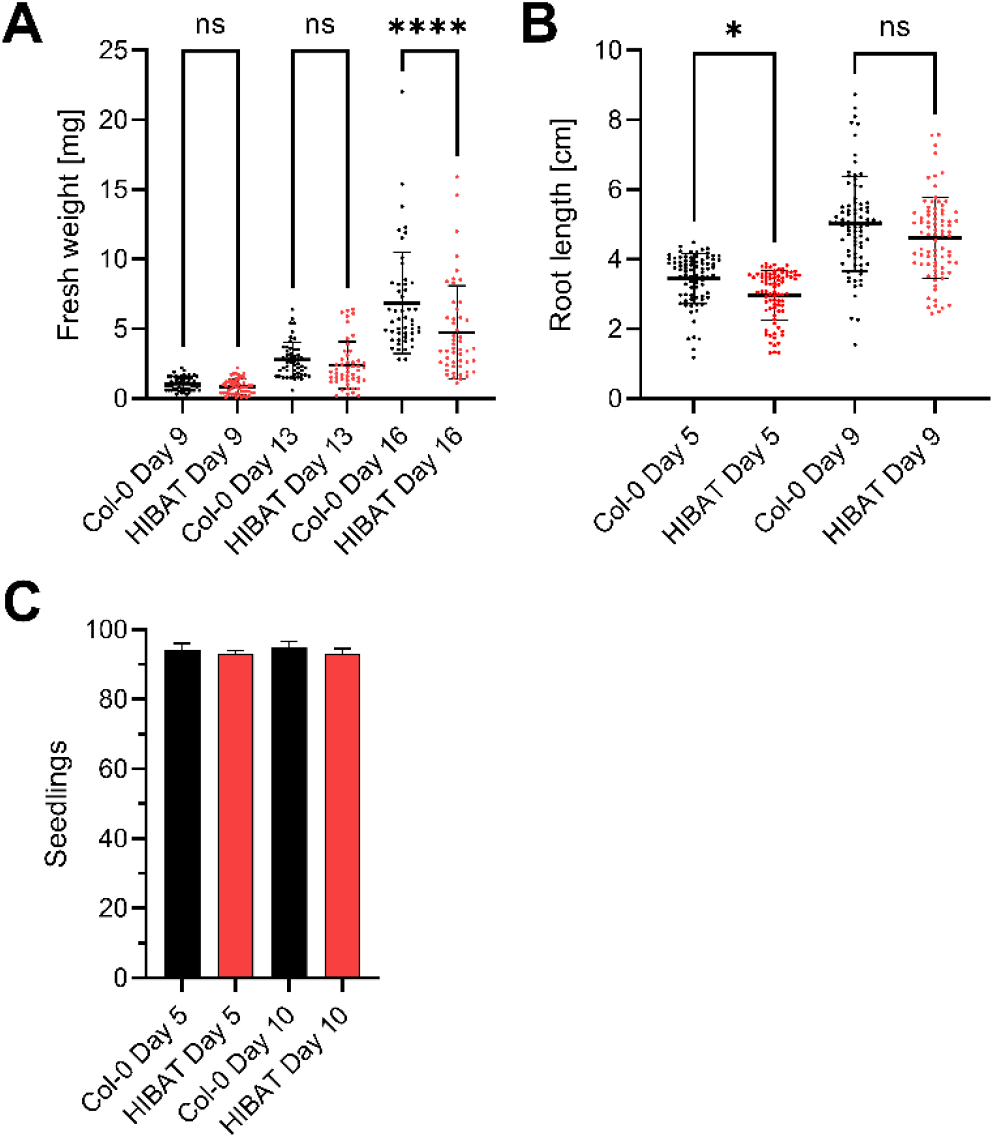
Physiology assays of HIBAT compared to Col-0. A) Fresh weight from shoots at 9, 13, and 16 days old of HIBAT and Col-0 lines (50 seedlings). B) Root length at 5 and 10 days old of both HIBAT and Col-0 plants (80 seedlings). C) Percentage of germination of the seed in HIBAT and Col-0 lines at 5 and 10 days old (100 seedlings). Asterisks indicate statistically significant differences determined by 2-way ANOVA (* P<0.05, ** P<0.01, *** P<0.001, **** P<0.0001, NS, Not Significant).

**Figure S 4 :**
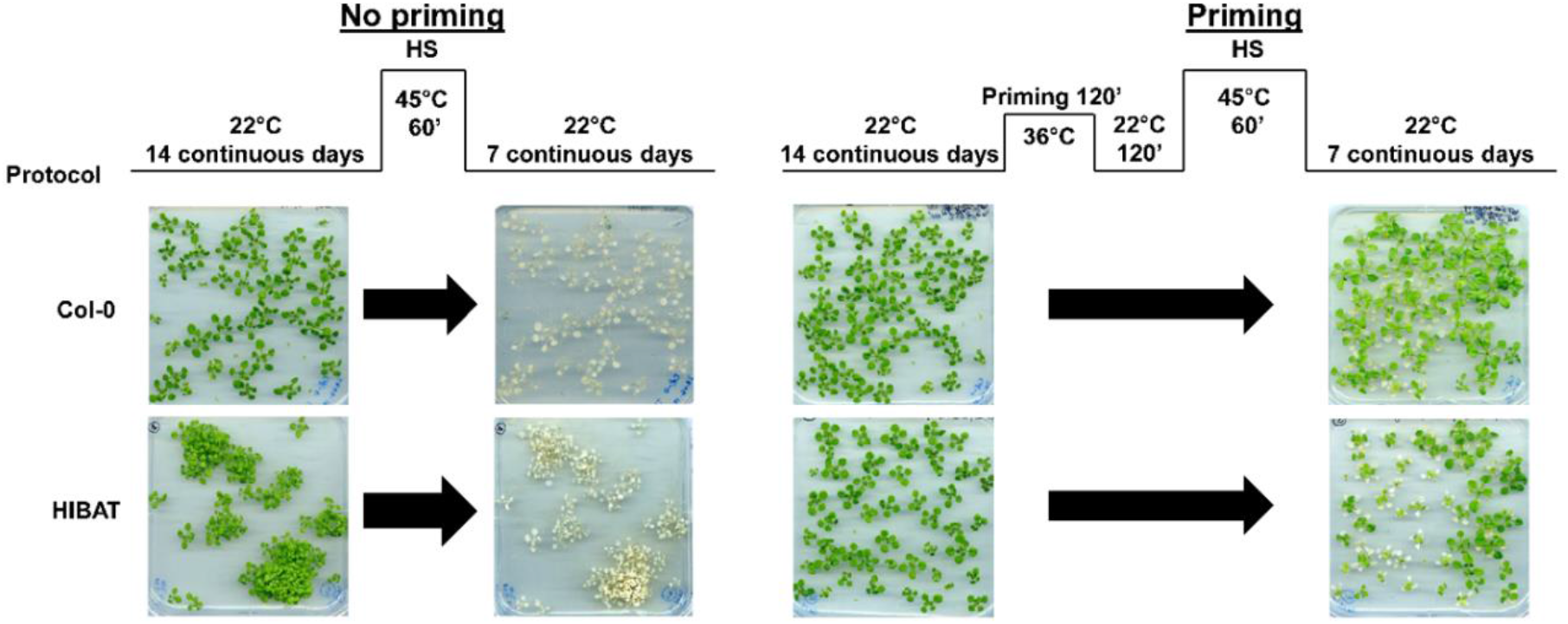
Acquired thermotolerance assay in the HIBAT and Col-0 lines. Left: no priming, right: with priming. Conditions applied for each plant are represented on top (protocol). Pictures were imaged at 14 and 21 days old. Percentage of dead cotyledons but surviving plants: around 46% in Col-0 and 76% in HIBAT.

**Figure S 5 :**
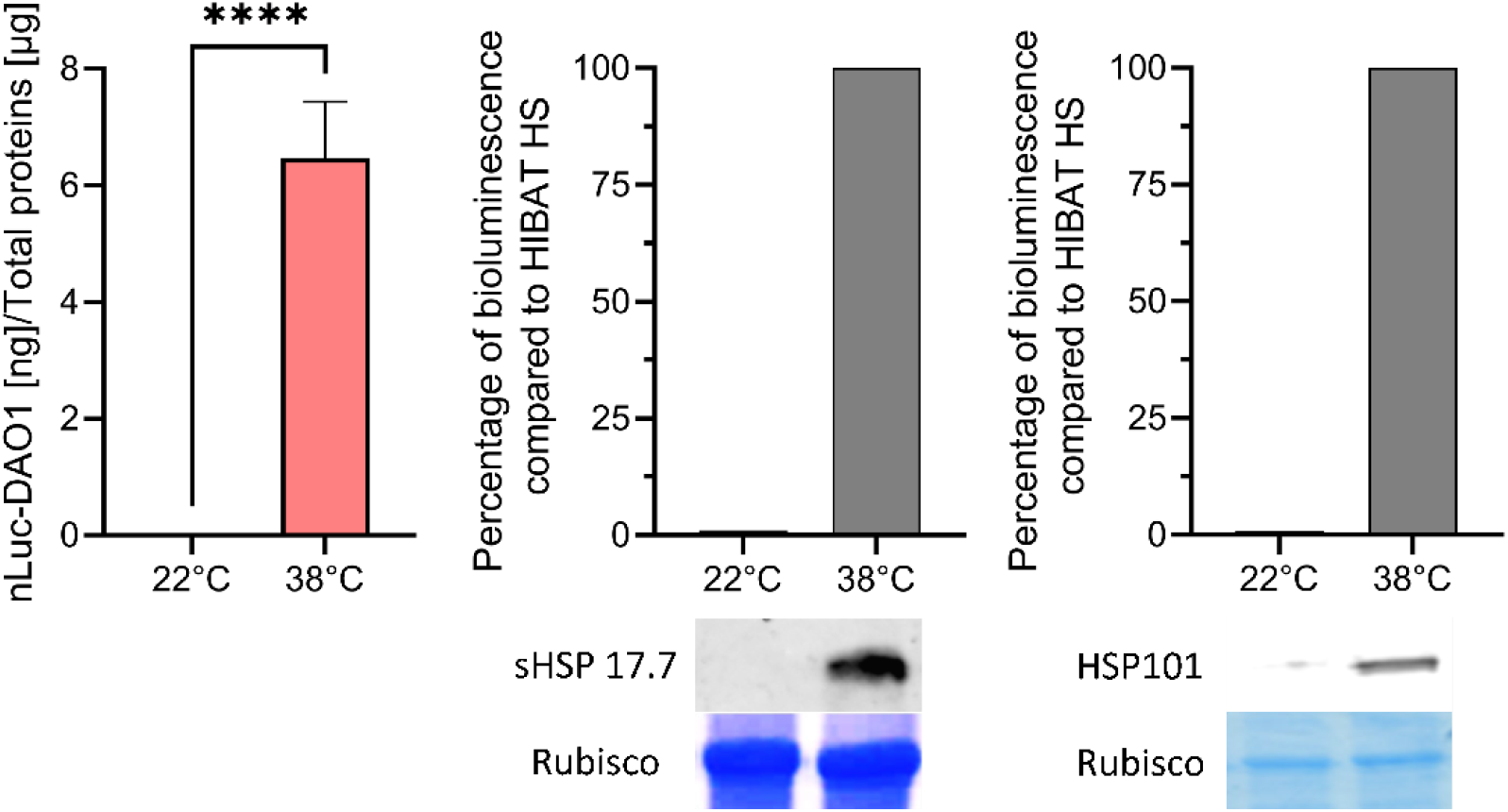
Left: Amounts of accumulated nLUC-DAO in 2 weeks old HIBAT seedlings, pretreated for 1 hours at either 22°C or 38°C, deduced from the relative nanoluciferase activity in crude extracts. Bellow: Immunodetection of Arabidopsis HSP17.7 (Right) and HSP101 (Left) in HIBAT seedlings pretreated, or not, for one hour at 38°C.

**Figure S 6 :**
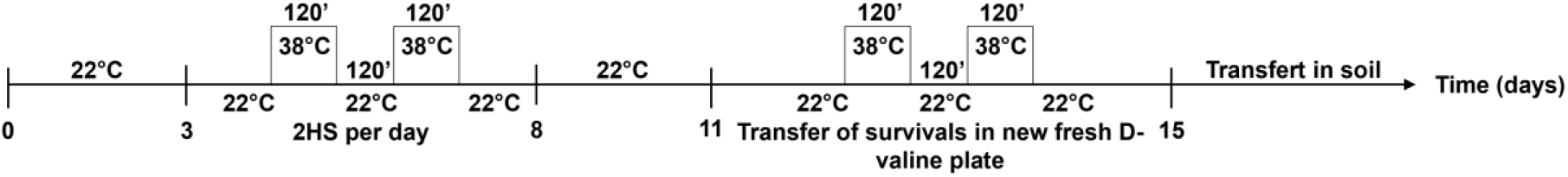
Experimental design for iterative heat stress.

**Figure S 7 :**
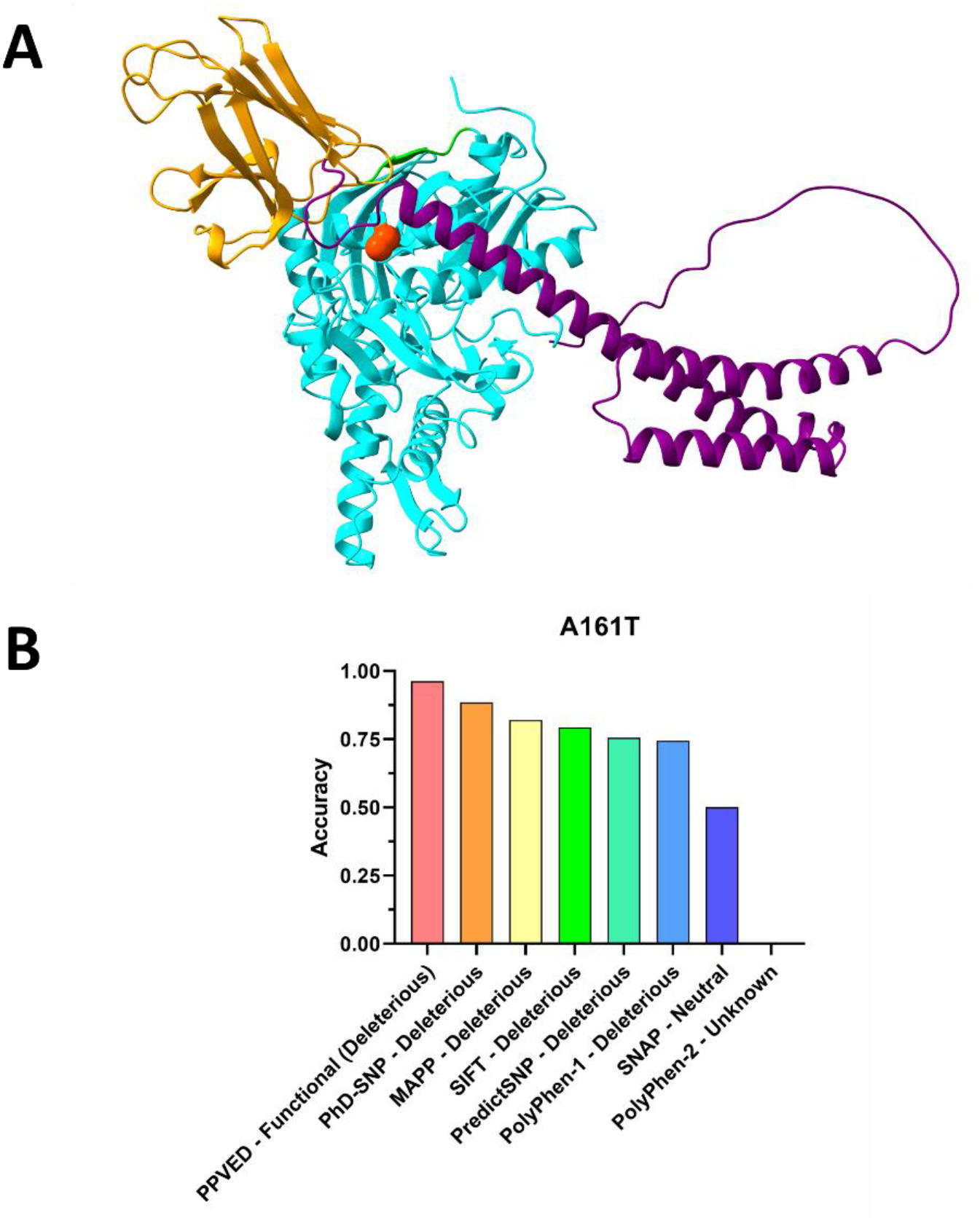
Effect of the point mutation A161T in HSP70-3. **A**) AlphaFold structure for Arabidopsis HSP70-3 (in the apo or ATP-bound state). Blue: nucleotide binding domain (NBD). Green: inter domain linker. Orange: substrate binding domain base (SBDα) Purple: Substrate biding domain lid (SBDβ). Red: Alanine 161 belongs to an alpha helix of the NBD, in a pocket facing the lid of the SBD. The lateral chain of threonine occupies of greater volume, and even more so when it is phosphorylated. **B**) Results of the analysis of different SNP detection tools on the Alanine to Threonine mutation in the alpha helix of HSP70-3. In Y, the accuracy of the measurement and in X the name of the program used and the effect of the mutation. Neutral: no difference; deleterious: negative effect on the protein; Unknown: the program is not able to detect a result. For PPVED, a so-called functional effect is one that affects the stability of the protein, PPVED is a plant specific tool.

## Notes

### Competing Interest Statement

The authors have declared no competing interest.

